# *In-silico* identification of Tyr232 in AMPKα2 as a dephosphorylation site for the protein tyrosine phosphatase PTP-PEST

**DOI:** 10.1101/2022.05.09.491063

**Authors:** Amrutha Manikandan, T S Sreevidya, Narayanan Manoj, Satyavani Vemparala, Madhulika Dixit

## Abstract

The AMP-activated protein kinase (AMPK) is known to be activated by the protein tyrosine phosphatase non-receptor type 12 (PTP-PEST) under hypoxic conditions. This activation is mediated by tyrosine dephosphorylation of the AMPKα subunit. However, the identity of the phosphotyrosine residues remains unknown. In this study we first predicted the structure of the complex of the AMPKα2 subunit and PTP-PEST catalytic domain using bioinformatics tools and further confirm the stability of the complex using molecular dynamics simulations. Evaluation of the protein-protein interfaces indicates that residue Tyr232 is the most likely site of dephosphorylation on AMPKα2. In addition, we explored the effect of phosphorylation of PTP-PEST residue Tyr64 on the stability of the complex. The phosphorylation of Tyr64, an interface residue, enhances the stability of the complex via the rearrangement of a network of electrostatic interactions in conjunction with conformational changes in the catalytic WPD loop. Our findings present a plausible structural basis of AMPK regulation mediated by PTP-PEST and shows how phosphorylation of PTP-PEST could be involved in its activation.

## 1 Introduction

Protein tyrosine phosphatase non-receptor type 12 (PTP-PEST) is a ubiquitously expressed cytosolic tyrosine phosphatase essential for embryonic development^1^. It mediates key cellular processes such as cell adhesion, migration and spreading, apart from being an important player in cancer^2–5^. PTP-PEST is 780 amino acids long and is composed of an N-terminal catalytic domain and a C-terminal domain containing four PEST (Proline, Glutamate, Serine and Threonine - rich) motifs^6^. The catalytic domain harbours the conserved ‘HC(X)5R’ motif responsible for the tyrosine phosphatase activity, while, the C-terminal domain is predominantly involved in binding with its substrates. The catalytic activity of PTP-PEST is significantly modulated by the phosphorylation of a conserved Y64 (Supplementary Figure S1) located in the active site^8^. Y64 forms aromatic stacking interactions with the sidechains of phosphotyrosines in several PTP-phospho-peptide substrate complexes^15–18^. As of date, over 18 substrates of PTP-PEST have been identified, which include MET, PDGFRβ, Cas, Paxillin, FAK, Pyk2 and HER2, among others^5,7^. The crystal structure of the catalytic domain of human PTP-PEST revealed the active site structural features that are the key determinants of both catalysis and substrate specificity^8^. Additionally, crucial residues and structurally plastic loop regions that enable PTP-PEST to recognize the substrate HER2 phosphorylation sites have been identified (Supplementary Figure S1)^8^.

Using cellular and enzymatic approaches, we recently demonstrated for the first time that the catalytic domain of PTP-PEST interacts with the α subunits of AMPK (5’-adenosine monophosphate activated protein kinase) to mediate the tyrosine dephosphorylation and consequent activation of the latter^9^. AMPK is a serine/threonine kinase that plays an important role in cell metabolism, cellular stress response and autophagy^10^. It is a heterotrimeric enzyme consisting of α, β and γ subunits, with each subunit having different isoforms (two α, two β and three γ isoforms)^11^. The AMPKα subunit that interacts with PTP-PEST contains the kinase domain (KD) as well as the autoinhibitory domain (AID)^9^. Evidence is available for multiple modes of regulation of AMPK kinase activity, including the phosphorylation of Y436, which negatively regulates its enzyme activity^12,13^. Our study established that both isoforms of AMPKα (α1 and α2) exist in complex with PTP-PEST under normoxic conditions and are dephosphorylated by PTP-PEST in response to hypoxia^9^. Specifically, AMPKα2 has been implicated in several hypoxia signalling pathways^14^. However, the identity of the potential AMPKα2 tyrosine residue(s) and the structural basis of how PTP-PEST recognizes and dephosphorylates AMPKα2 remains elusive. Given the lack of the crystal structure of a substrate-bound complex of PTP-PEST, here, we use *in-silico* approaches to identify Y232 as a potential tyrosine dephosphorylation site in the KD of AMPKα2. Furthermore, using protein-protein docking in combination with molecular simulations we propose the structural basis of the dephosphorylation of AMPKα2 by PTP-PEST and show how this activity is modulated by the phosphorylation status of residue Y64 in PTP-PEST. The results are consistent with the notion that AMPK is indeed a new physiological substrate of PTP-PEST.

## 2 Materials and methods

### 2.1 Prediction and selection of PTP-PEST dephosphorylation sites on the AMPKα2 subunit

In order to identify the potential tyrosine dephosphorylation sites on AMPK, the amino acid sequence of human AMPKα2 (gene *PRKAA2*, gene ID: 5563) was used to predict the tyrosine phosphorylation sites using the NetPhos 3.1^20^ and GPS 3.02^21^ web servers. A threshold of phosphorylation potential greater than 0.6 was applied to the NetPhos3.0 predictions and the highest threshold was used for GPS3.0 predictions. Putative sites that were common between both predictions were chosen for a further assessment of whether they could be dephosphorylated by PTP-PEST. The sequence was next examined in the PhosphoSitePlus^22^ database for experimentally reported evidence on phosphorylation.

### 2.2 Phosphorylation and structure preparation

The crystal structure of the catalytic domain of human PTP-PEST (PDB ID: 5HDE) was retrieved from the Protein Data Bank (PDB) and prepared using UCSF Chimera^23^. The modified catalytic residue, CSP231 (phosphorylated cysteine) was replaced with cysteine. The crystal structure of the isolated KD of inactive AMPKα2 containing residues, 10-278 (PDB ID: 2H6D) was retrieved from PDB and two missing regions, namely residues 167 to 179 and 279-288 were modelled using MODELLER^24^ utilizing the structure of the full-length protein as the reference (PDB ID: 5ISO). The resulting structure was corrected for clashes and energy minimized using steepest descent for 1000 steps. Next, the predicted tyrosines were individually phosphorylated using UCSF Chimera^23^.

### 2.3 Molecular docking analysis

Experimental data^8, 25^ on PTP-PEST was used to guide docking of the PTP-PEST catalytic domain and AMPKα2 KD (henceforth referred to as PTP-PEST and AMPK respectively) using the ClusPro 2.0^26^ program. This program performs rigid-body docking using PIPER, which is a correlation approach based on Fast Fourier Transform (FFT). ClusPro has consistently performed well in the Critical Assessment of Prediction of Interactions (CAPRI) and its use of structure-based pairwise interaction along with energy functions provides near native models after docking. We refined the docking using surface accessible PTP-PEST active site residues Y64, D66, R140, K142, H200, H274 and Q278 as “attraction residues”^8^. Previous site directed mutagenesis studies and molecular dynamics analyses have revealed the role of these residues in catalytic activity or substrate recognition by PTP-PEST^8,26^. Similarly, the residues corresponding to the predicted phosphotyrosine and the neighbouring pY-1, pY+1 residues on AMPKα2 were also denoted as “attraction residues”, with all other docking parameters set to default. PTP-PEST was set as the receptor and AMPKα2 KD as ligand. We used “balanced” energy coefficients and evaluated the largest cluster from each of the docking experiments. The top ten complexes from the largest cluster in each experiment were repaired using the RepairPDB function of the FoldX^27^ plugin in YASARA^28^.

### 2.4 Assessment of docked structures and interfaces

The top 10 models obtained after docking PTP-PEST with each phosphorylated AMPK model were evaluated for their binding free energies and dissociation constants (Kd) predicted using the PRODIGY^29^ program. In PRODIGY, the IC value is defined as the number of interfacial contacts between the two interacting proteins, which are classified according to the nature of these residues as polar, apolar or charged. Two residues are defined in contact if any two of their heavy atoms are within a distance of 5.5 Å. It also considers the properties of non-interacting surfaces (NIS) and uses the following equation^29^:

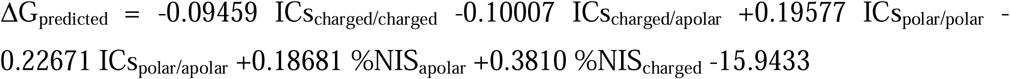

The binding affinity of PTP-PEST with each potential phosphotyrosine on the AMPK was further evaluated using the PRODIGY-LIGAND program ^29^. This uses the atomic contacts (ACs) within the distance threshold of 10.5 Å and classifies the ACs according to the atom involved in the interaction (C=Carbon, O=Oxygen, N=Nitrogen, X=all other atoms), using the following equation^29^:

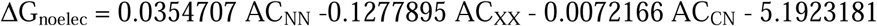

The interfaces of all complexes were also evaluated for their stability using PDBePISA^30^. Two parameters Δ^i^G, indicating the solvation free energy gain upon formation of the interface as well as the Δ^i^G P-value, which is an indicator of an interaction-specific interface-at a value < 0.5, were examined. Based on these parameters and manual examination, one docked PTP-PEST—AMPK complex was chosen as a representative model for the interaction between PTP-PEST and AMPK phosphorylated separately at a specific tyrosine. The representative model was then evaluated for the protein-protein interaction energy using the FoldX^27^ program. This function calculates the interaction energy by first unfolding the target proteins, then subtracting the sum of the individual energies from the global energy.

### 2.5 Molecular dynamics simulations

The stability of the chosen PTP-PEST—AMPK(pY232) complex was examined *via* all-atom classical molecular dynamics simulations of the complex by using the CHARMM36 forcefield^31^ with NAMD software^32^. The correct protonation states of the ionizable residues in the complex were determined at a pH of 7.2 using DEPTH^33^ server and appropriately changed before solvating the system with the TIP3P water system^34^. To maintain overall charge neutrality and to mimic physiological salt concentration, 0.15 mol/L NaCl was added to each system. The docked complex was first energy minimized to reduce any steric clashes which may be present after docking. The complex was then equilibrated for 100ns in the isothermal-isobaric (pressure is 1 atm and temperature is 300K) ensemble. The resultant trajectory was analysed to check for stability of the complex via monitoring the root mean squared deviation from the docked structure. After ascertaining the stability of the docked complex, the next step was to understand the effect of phosphorylating Y64, a key residue of PTP-PEST on the docked complex. Towards this goal, we simulated two systems, PTP-PEST—AMPK(pY232) (referred to as control) and PTP-PEST(pY64)—AMPK(pY232) (referred to as pY64, where pY stands for PTP-PEST phosphorylated at Y64), for an additional 200 ns each. The structure at the end of the 100ns equilibration run was taken as the starting point for the two simulations. The details of the systems studied are shown in Table 1. VMD software^35^ and the PyMOL Molecular Graphics System, Version 2.4.2 Schrödinger, LLC^36^ were used for visualization and analysis was done using in-house analysis codes using TCL scripting language and VMD plugins.

**Table 1.** Selection of a representative model from the docked PTP-PEST—AMPK complexes

| **A. PTP-PEST—AMPK(pY104)** |  |  |  |  |  |
| --- | --- | --- | --- | --- | --- |
| **Model No.** | **$\Delta G$ PRODIGY**<br>**(Kcal/mol)** | **PRODIGY Kd**<br>**(M)** | **$\Delta G$ PRODIGY LIGAND**<br>**(Kcal/mol)** | **$\Delta^i G$ (Kcal/mol)** | **$\Delta^i G$ P-**<br>**value** |
| 0 | -12.4 | 8.30E-10 | - | -6.5 | 0.446 |
| 1 | -11.6 | 3.00E-09 | -6.7 | -1.2 | 0.621 |
| 2 | -13.9 | 6.90E-11 | -6 | -6.4 | 0.557 |
| 3 | -12.4 | 5.10E-10 | - | -3.6 | 0.657 |
| 4 | -13.7 | 8.80E-11 | -6.5 | -8 | 0.264 |
| 5 | -17.1 | 2.90E-13 | -5.9 | -9.6 | 0.477 |
| 6 | -13.4 | 1.50E-10 | -6.6 | -5.8 | 0.569 |
| 7 | -11.7 | 2.80E-09 | -6 | -5.4 | 0.632 |
| 8 | -12.9 | 3.30E-10 | -5.8 | -2 | 0.737 |
| 9 | -11.1 | 7.60E-09 | -6 | -2.8 | 0.665 |
| **B. PTP-PEST—AMPK(pY283)** |  |  |  |  |  |
| **Model No.** | **$\Delta G$ PRODIGY**<br>**(Kcal/mol)** | **PRODIGY**<br>**Kd (M)** | **$\Delta G$ PRODIGY LIGAND**<br>**(Kcal/mol)** | **$\Delta^i G$ (Kcal/mol)** | **$\Delta^i G$ P-**<br>**value** |
| 0 | -8.6 | 4.70E-07 | -6.4 | -2.8 | 0.67 |
| 1 | -10.6 | 1.60E-08 | -6.1 | 0.4 | 0.863 |
| 2 | -9 | 2.40E-07 | -6.1 | -2.1 | 0.713 |
| 3 | -8.7 | 4.40E-07 | -6.4 | -2.7 | 0.825 |
| 4 | -12.1 | 7.70E-10 | - | -9.5 | 0.461 |
| 5 | -8.7 | 3.90E-07 | -5.8 | -2.4 | 0.657 |
| 6 | -9.1 | 2.20E-07 | -6 | -1.8 | 0.809 |
| 7 | -11.3 | 5.10E-09 | -6.3 | -3.5 | 0.716 |
| 8 | -9.3 | 1.40E-07 | -6 | 0.5 | 0.833 |
| 9 | -12.9 | 3.50E-10 | -6 | 2.2 | 0.9 |
| **C. PTP-PEST—AMPK (pY232)** |  |  |  |  |  |
| **Model No.** | **$\Delta G$ PRODIGY**<br>**(Kcal/mol)** | **PRODIGY**<br>**Kd (M)** | **$\Delta G$ PRODIGY LIGAND**<br>**(Kcal/mol)** | **$\Delta^i G$ (Kcal/mol)** | **$\Delta^i G$ P-**<br>**value** |
| 0 | -9.1 | 2.00E-07 | -6.5 | -5.2 | 0.537 |
| 1 | -12.6 | 5.40E-10 | -7.1 | -3.7 | 0.636 |
| 2 | -9.9 | 5.50E-08 | -6.7 | -1.1 | 0.514 |
| 3 | -14.7 | 1.60E-11 | -7.3 | -9.7 | 0.4 |
| 4 | -11.1 | 7.30E-09 | -6.3 | -5.9 | 0.571 |
| 5 | -10.3 | 2.90E-08 | -6.8 | -10.5 | 0.259 |
| 6 | -10.6 | 1.60E-08 | -6.4 | -3.7 | 0.672 |
| 7 | -11.6 | 2.90E-09 | -7.1 | -4.7 | 0.555 |
| 8 | -11.1 | 7.10E-09 | -6.8 | -8.4 | 0.376 |
| 9 | -11.8 | 2.10E-09 | -6.7 | -7.8 | 0.446 |

\*The '-' value for $\Delta G$ PRODIGY LIGAND (Kcal/mol) scores indicates that there was no interaction of the phosphotyrosine with PTP-PEST residues

### 2.6 Community network analysis

Community network analysis is a tool to identify connections between amino acids of the protein based on their classification into communities that are derived from the molecular dynamics trajectories^37^. The community network analysis on the MD trajectory data was done using NetworkView plugin in VMD^38^. A network is a set of nodes connected using edges. A node is defined as each C*α* atom of amino acid in the protein. Edges connect pairs of nodes if the Cα atoms of the corresponding residues are within 4.5□Å of each other for at least 75 % of the frames analyzed. In this analysis, edges between Cα atoms that have adjacent residue numbers are disallowed. The edges are weighted using correlation matrix (C*ij*) data between the C*α* atoms using the relation;

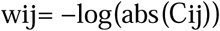

These weights in the form of correlation matrices are calculated using Carma software ^39^. The correlations from residue motions are used as a measure for information transfer between the two residues in contact. The community detection analysis was done using software “gncommunities”^38^. Community network analysis was done on control-PEST-AMPK (control) and pY64-PEST—AMPK (pY64) complexes in VMD using the NetworkView plugin in VMD.

### 2.7 Free energy calculations

MM-GBSA (Molecular Mechanics-Generalised Born Surface Area) calculations ^41–43^ was performed to calculate the free energy of the protein complex systems.

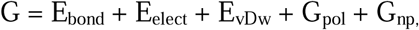

where the first three terms are bonded (bond, angle and dihedral), electrostatic and van der Waals interactions calculated from the MD trajectories. *G*_pol_ and *G*_np_ are the polar and non-polar contributions to the solvation free energies. *G*_pol_ is typically obtained by solving the generalized Born (GB) model (given the MM/GBSA approach), whereas the non-polar term is estimated from a linear relation to the solvent accessible surface area (SASA).

## 3 Results

### 3.1 Putative PTP-PEST dephosphorylation sites are localized in the AMPKα2 kinase domain

The NetPhos and PhosphoSitePlus programs predicted seven potential tyrosine dephosphorylation sites on full-length AMPK (Figure 1A, Supplementary Table S1). We further analysed the primary sequence neighbouring these sites to assess their viability for dephosphorylation by PTP-PEST. Although consensus motifs for substrates have not been reported for PTP-PEST or other protein tyrosine phosphatases (PTPs), it is known that PTPs have an intrinsic sequence selectivity and recognise at least 3-5 residues on either side of the pY (phosphotyrosine)^24^. Studies by Selner et al. show that PTP-PEST exhibits a strong affinity towards acidic residues and disfavours basic residues that immediately precede/follow the phosphorylated tyrosine^24^. Furthermore, using HER2 substrate peptides, Li et al. demonstrated that PTP-PEST strongly favours peptides with acidic residues at the pY-1 and pY-4 positions, while Selner et al., used synthetic peptide libraries to establish that PTP-PEST prefers hydrophobic residues at the pY-1 position and exhibits a C-terminal selectivity for acidic residues^8,24^. The seven selected sites were evaluated based on the presence of acidic or hydrophobic residues at the pY-1 and pY+1 positions and the absence of basic residues within the pY+2 and pY-2 positions (Figure 1B). This assessment helped identify the three most suitable tyrosine phosphorylation sites on AMPK, namely Y104, Y232 and Y283 (Figure 1C and Figure 1D), present in the AMPKα2 subunit. Examination of sequence conservation of these three residues showed that Y104 is largely conserved across AMPKα2 in vertebrates and its homolog (Snf1) in fungi (Figure 1E, Supplementary Figure S2 A). However, in Snf1, the residue corresponding to Y232 is substituted (often by threonine) and the region corresponding to Y283 is deleted (Figure 1E, Supplementary Figure S2 B and C). Given the absence of the PTP-PEST ortholog in fungi, it is evident that the mode of interaction of the AMPKα2 and PTP-PEST as described in the following sections is specific to only those organisms that contain the corresponding pairs of orthologs.

**Figure 1.**
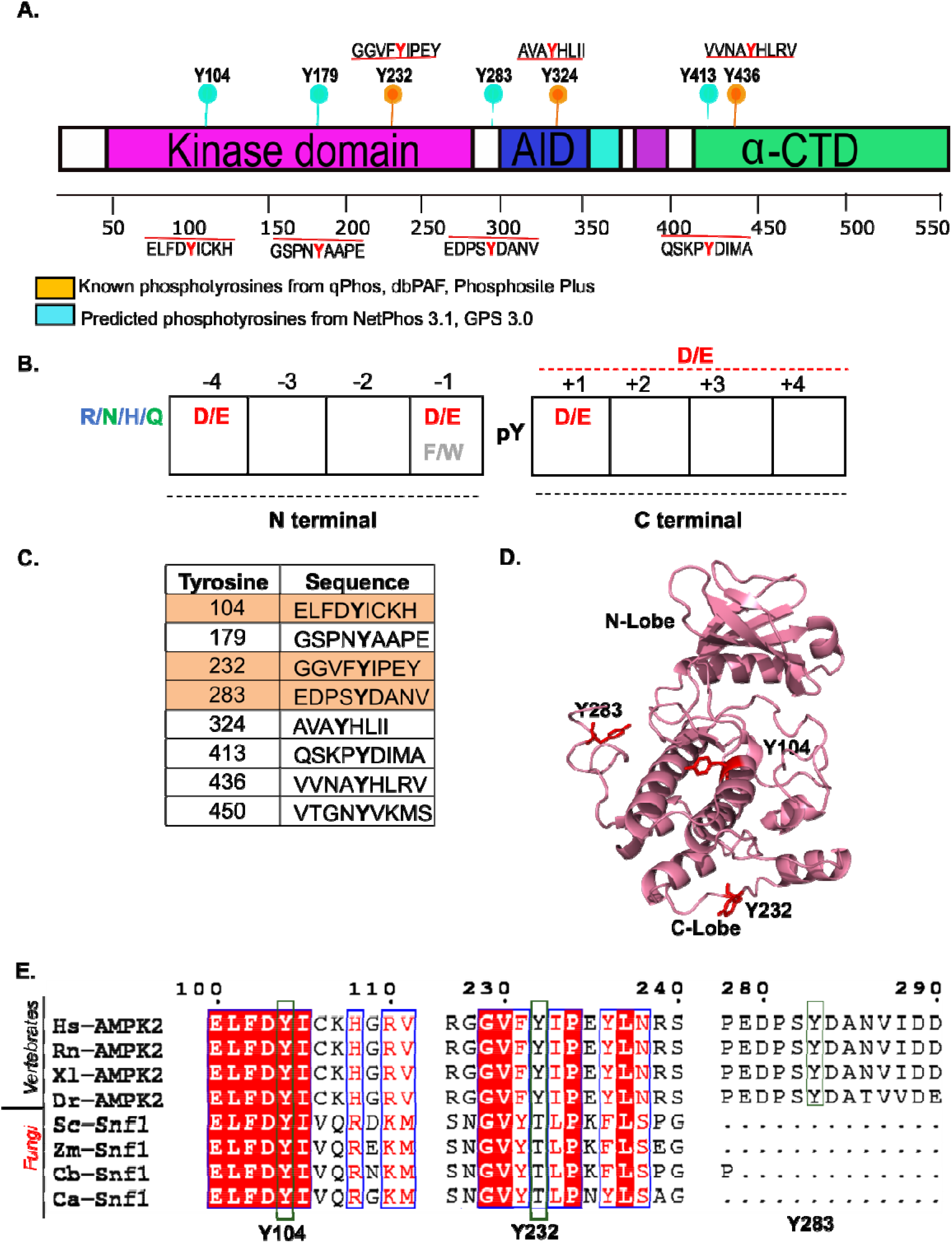
Identification of putative tyrosine phosphorylation sites on human AMPKα2. (A) Cartoon depicting predicted and known tyrosine phosphorylation sites on AMPKα2. (B) Sequence characteristics governing the phosphopeptide selectivity of PTP-PEST. (C) Primary peptide sequences of the putative phosphorylation sites on AMPKα2. (D) Ribbon representation of the kinase domain of AMPKα2 with the residues Y104, Y232 and Y283 depicted in stick representation. (E) Multiple sequence alignment of regions in AMPKα2 (vertebrates) and its fungal homolog, Snf1 (Y104, Y232 and Y283, are shown in black boxes). The alignment between *Homo sapiens* (Hs), *Rattus norevgicus* (Rn), *Xenopus laevis* (Xl), Danio rerio (Dr), *Saccharomyces cerevisae* (Sc), *Zygosaccharomyces mellis* (Zm), *Candia glabrata* (Cb) and *Candida albicans* (Ca), are represented in this image.

### 3.2 Docking analysis of PTP-PEST with phosphorylated AMPK

Our next goal was to model the structure of the complex of PTP-PEST and AMPK. Towards this, three docked complex structures of PTP-PEST with AMPK individually phosphorylated at the predicted tyrosine phosphosites (pY104, pY232 and pY283, where pY refers to phosphotyrosine) were constructed using ClusPro^25^. The docked structures were then evaluated for their suitability to represent the best possible binding mode of PTP-PEST with AMPK. The highest ranked cluster (Supplementary Tables S2-4) from each docking run was chosen and the top 10 models of that cluster were examined. We first analysed each model manually using the primary criterion that the phosphotyrosine is located at the protein-protein interface. Each complex was then assessed for its protein-protein binding affinity (Kcal/mol) and for the free energy of ligand binding i.e. the binding affinity of PTP-PEST with the phosphorylated tyrosine on the AMPK, followed by evaluation of their interface using PDBePISA^30^. Based on these analyses, a single model was chosen as the best model of a representative complex for each of the three predicted phosphosites (Table 1).

For the complex PTP-PEST—AMPK(pY104), Model 4 was chosen as a representative for its highest values of favourable protein-protein binding affinity (−13.7 Kcal/mol) and energy of interface formation (Table 1A). Model 4 showed a Δ^i^G P-value of 0.264, indicating higher hydrophobicity than that expected for a non-specific interface, and therefore, a structurally significant association. The phosphorylated Y104 interacts with the active site cleft residues R43, S275 and R270 of PTP-PEST (Figure 2A and Figure 2B, Supplementary Figure S3). However, pY104 does not directly interact with either of the catalytic residues, C231 and R237 on the P-loop of PTP-PEST (Figure 2B, Supplementary Figure S3). Similarly, Model 7 was chosen as a representative for the complex PTP-PEST—AMPK(pY283) for its most favourable protein-protein binding affinity and energy values (Table 1B). However, in all these models, neither pY283 (Figure 2C and Figure 2D) nor pY104 displayed direct interactions with the catalytic P-loop residues of PTP-PEST (Figure 2, Supplementary Figure S3 and S4).

**Figure 2.**
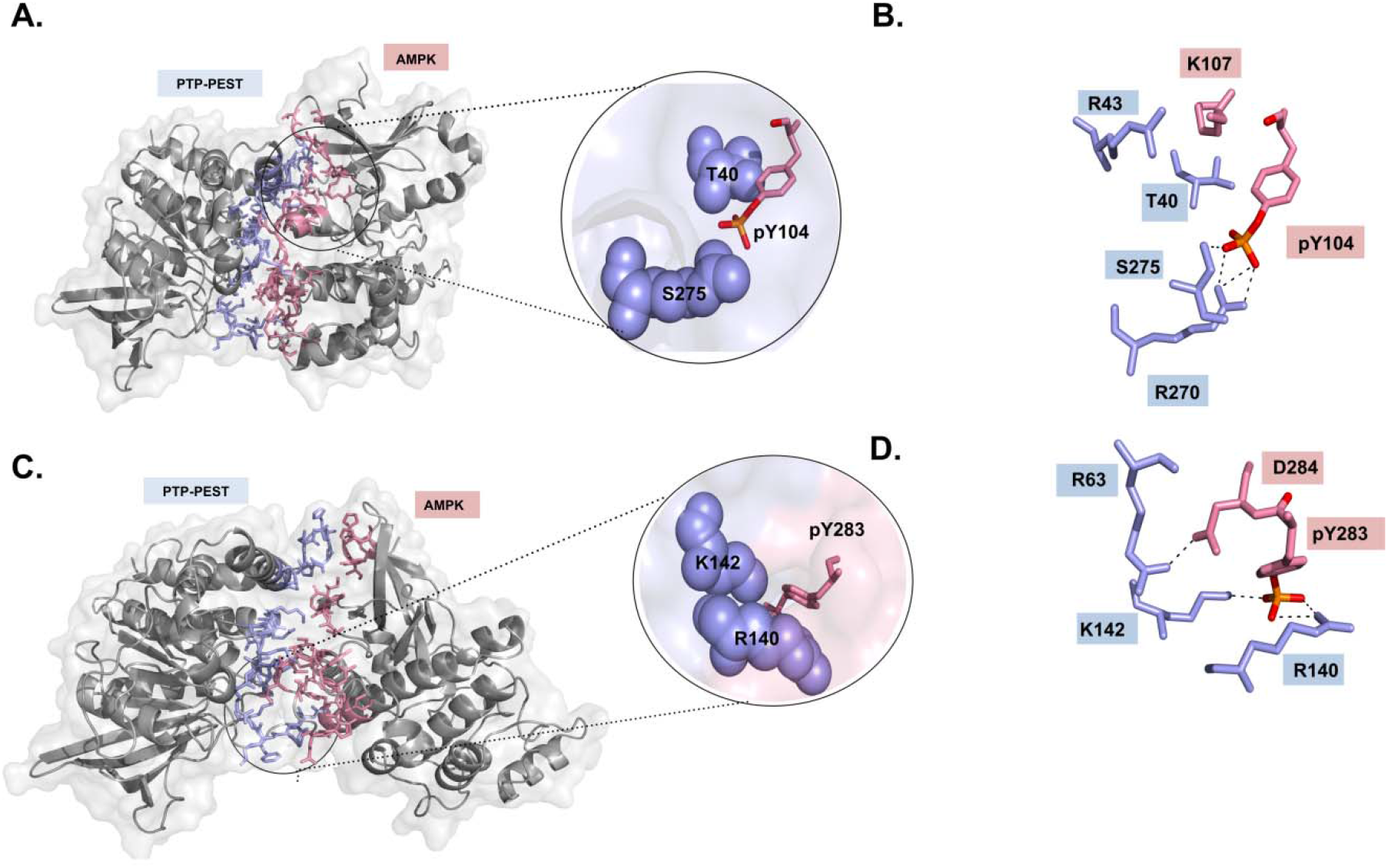
Docked complexes of PTP-PEST—AMPK(pY104) and PTP-PEST—AMPK(pY283). (A) Surface representation of the docked complex showing the interactions of pY104 on AMPK (pink) with a cleft formed by the residues T40 and R270 of PTP-PEST (sky-blue). (B) Stick representation of interactions between residue pY104 in AMPK and those of PTP-PEST. (C) Surface representation of the docked complex showing the superficial interactions of pY283 on AMPK (pink) with a cleft formed by the residues R140 and K142 of PTP-PEST (sky-blue). (D) Stick representation of the interface interactions between pY283 on AMPK and residues on PTP-PEST. The dashed lines represent non-covalent interactions (≤3.5Å), between residues.

For the PTP-PEST—AMPK(pY232) complex, Model 3 was chosen as the representative. Model 3 displayed both a high negative value of protein-protein binding affinity (−14.7 Kcal/mol) as well as a Δ^i^G P-value of 0.4, indicating a stable complex between PTP-PEST and AMPK, with a structurally significant interface (Table 1C). Moreover, this model showed the highest negative ligand binding energy between PTP-PEST and AMPK(pY232), among all models considered for the three phosphosites (Table 1). Interestingly, unlike the two PTP-PEST—AMPK(pY104) and PTP-PEST— AMPK(pY283) complexes, in this case, the phosphotyrosine is positioned in the catalytic site of PTP-PEST, within hydrogen bonding distance of the catalytic residues C231 and R237 (Figure 3A,B and Supplementary Figure S5). Given these observations, it appears that the PTP-PEST—AMPK(pY232) complex is the most plausible model of an enzyme-substrate complex where pY232 is dephosphorylated by PTP-PEST. Notably, pY232 interacts with PTP-PEST residues, namely, H200, which is known to stabilize the incoming phosphotyrosines and with the catalytic proton donor D199 (Figure 3B). The interface of this docked complex was also populated with other critical residues, namely Y64, H274 and K280, which are known to be involved in substrate stabilization (Figure 3B, Supplementary Figure S5). On comparison of the three complexes, we also observed that the PTP-PEST—AMPK(pY232) complex showed a greater number of interface contacts compared to the other two (Supplementary Table S5). Additionally, the FoldX interaction energy, which is a consolidation of all the energies required to form a complex, was lowest (most favourable) in the PTP-PEST—AMPK(pY232) complex (Supplementary Table S5).

**Figure 3.**
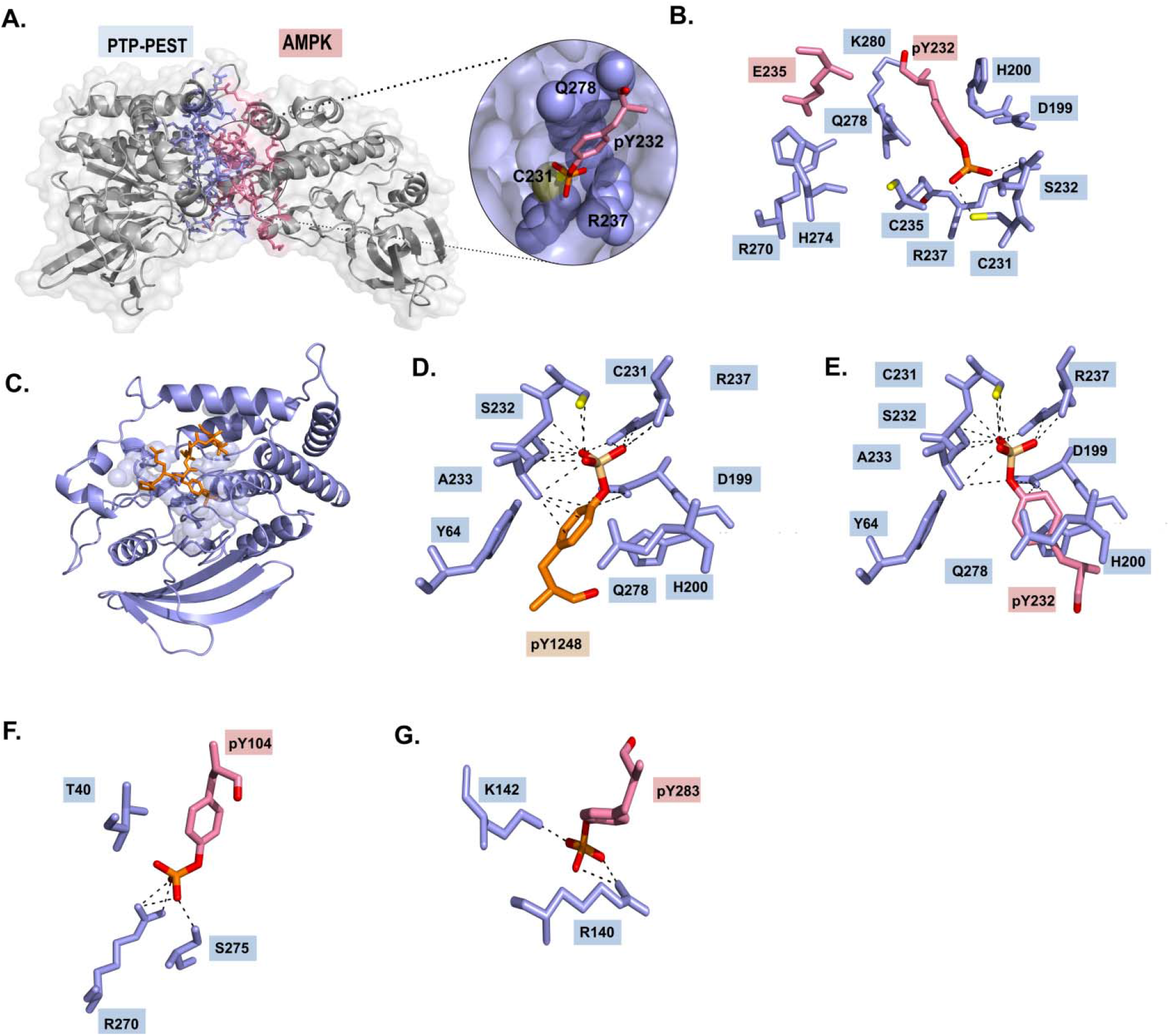
Comparison of docked PTP-PEST—AMPK(pY232) complex and the model of PTP-PEST—HER2(pY1248) complex. (A) Surface representation of the docked complex showing the insertion of AMPK residue pY232 (pink) within a cleft formed by the residues C231, Q278, G236 and R237 of PTP-PEST. (B) Stick representation of the interface interactions between pY232 of AMPK and residues on PTP-PEST. (C) Ribbon representation of the model of PTP-PEST in complex with HER2(pY1248) heptapeptide. Closeup views in stick representations of the interactions between (D) pY1248 of HER2 heptapeptide and PTP-PEST, (E) pY104 of AMPK and PTP-PEST, (F) pY232 of AMPK and PTP-PEST and (G) pY283 of AMPK and PTP-PEST. The dashed lines represent non-covalent interactions (≤3.5Å) between two amino acids.

### 3.3 Comparison of the PTP-PEST—AMPK(pY232) complex with PTP-PEST HER2(pY1248) complex

Given that pY232 is the most likely AMPK phosphosite that is targeted by PTP-PEST, we attempted to establish the validity of the active site interactions of the PTP-PEST— AMPK(pY232) complex model. As of date, the sole crystal structure available of PTP-PEST is that of the apo form. Therefore, we used the crystal structure of the catalytic domain of PTPN18 complexed with a cognate HER2 phosphorylated heptapeptide (pY1248) (PDB ID: 4GFU), to create a model of the PTP-PEST—HER2-pY1248 peptide complex using superposition and manual positioning (Figure 3C). We believe that this model is a reasonable representation of a PTP-PEST—cognate phosphopeptide complex for the following reasons. First, both PTP-PEST and PTPN18 are homologous phosphatases and the catalytic domains share a high level of sequence and structural similarity with an RMSD of 1.0 Å over 261 Cα atoms (sequence identity of 47%). PTP-PEST and PTPN18 share a conserved active site with almost identical residues, although they do display a few differences, such as S229 in PTPN18 in place of the catalytic C231 in PTP-PEST (Supplementary Figure S6 A, B). Importantly, HER2 is a known substrate for both PTP-PEST and PTPN18 and both enzymes dephosphorylate HER2 at residue pY1248^8,15^. Lastly, the observed interactions between the active site residues and the modelled phosphopeptide is consistent with enzyme kinetic data of PTP-PEST wildtype and a series of active site mutants^26^. As expected, in the model, pY1248 interacts with the catalytic residues C231 and R237 (Figure 3D). Hence, we used this PTP-PEST— HER2(pY1248) complex as a reference to perform a comparative analysis with the three docked complexes of PTP-PEST and AMPK phosphorylated at Y104, Y232, and Y283. Comparisons showed that the interactions of pY232 with C231 and R237 of PTP-PEST are almost identical to those seen in the reference complex (Figure 3D and Figure 3E). However, in the case of the PTP-PEST—AMPK(pY104/pY283) complexes, there were no common contacts compared to the reference complex and the interactions do not reflect catalytically competent binding (Figure 3F and Figure 3G). It is to be noted that the PTP-PEST—AMPK(pY232) complex displays minor differences in interactions with respect to the reference complex. This is not entirely unexpected since binding modes of AMPK and HER2 to PTP-PEST may indeed be different or these arise because of the modelling process. Nevertheless, the above observations lend strong support for our prediction that pY232 is the most likely AMPK phosphosite that is regulated by PTP-PEST.

### 3.4 Effect of phosphorylation of PTP-PEST residue Y64 on the stability of the PTP-PEST—AMPK(pY232) complex

Y64 in PTP-PEST is a highly conserved residue that is present at the interface of the docked PTP-PEST—AMPK(pY232) complex (Supplementary Figure S5). Y64 is a critical residue known to be involved in substrate recognition. Substitution of the residue by alanine leads to a loss in the catalytic activity of PTP-PEST^8^. Moreover, Y64 is a known phosphorylation site of PTP-PEST, as documented in PhosphositePlus^21^. In order to examine the effect of phosphorylation of Y64 on the stability of the PTP-PEST— AMPK(pY232) complex, we carried out atomistic MD simulations and compared the relative stabilities of the PTP-PEST—AMPK(pY232) and the PTP-PEST(pY64)— AMPK(pY232) complexes (referred to as control and pY64 complexes, respectively). First, the control complex was subjected to a 100ns MD simulation during which we monitored the root mean squared deviation (RMSD), solvent accessible surface area (SASA) and radius of gyration (Rg) of the individual proteins in the complex (Supplementary Figure S7). The data suggests that AMPK undergoes more conformational changes as compared to PTP-PEST. These changes possibly reflect a response to the complex formation or phosphorylation of Y232 on AMPK. Alternatively, this could be because of flexibility introduced by a missing loop that was modelled in AMPK. Next, we used this complex and simulated two systems: with and without the phosphorylation of Y64 of PTP-PEST for 200ns each.

The overall stability of the two complexes was monitored *via* time evolution of RMSD and RMSF using the structure at the end of 100ns of simulation of docked complex (Figure 4), along with the solvent accessible surface area (SASA) and radius of gyration (Rg) (Supplementary Figure S8). Although both complexes are stable over the simulation time scales (Figure 4A and Figure 4B), a marked increase in the RMSD of AMPK in the initial phase of the simulation with Y64 phosphorylated suggests a conformational change (Figure 4A). This is also reflected in the changes in the RMSF values calculated over the last 25ns of simulation (Figure 4C and Figure 4D). The changes in mobility of the pY64 complex from RMSF data is highlighted in those regions showing marked changes compared to the control complex (Figure 4E). A significant number of residues showing mobility changes occur at the complex interface probably as an effect of phosphorylation of Y64. To further quantify the effect of pY64 on the energetics of the complex, we use MM-GBSA technique within NAMD software^32^ over the simulation trajectory (Figure 5A). The results are indicative of a predominant increase in attractive electrostatic interaction of the complex, where the change in energies is plotted (ΔE_pY64_-ΔE_control_) (Figure 5B). In addition, there is also an increase in the polar solvation energy of the complex. The electrostatic and van der Waals interaction energies (Vdw) between PTP-PEST and AMPK of the complex are also calculated over the simulation trajectory and show that in the pY64 complex, there is an increase in the favourable interactions between the two proteins. Taken together, the data strongly suggests that phosphorylation of Y64 leads to an enhanced binding affinity in the complex. We next looked at the origins of this variation in electrostatic interaction energy by probing the local interaction network at the complex interface and also within individual proteins of the complex.

**Figure 4.**
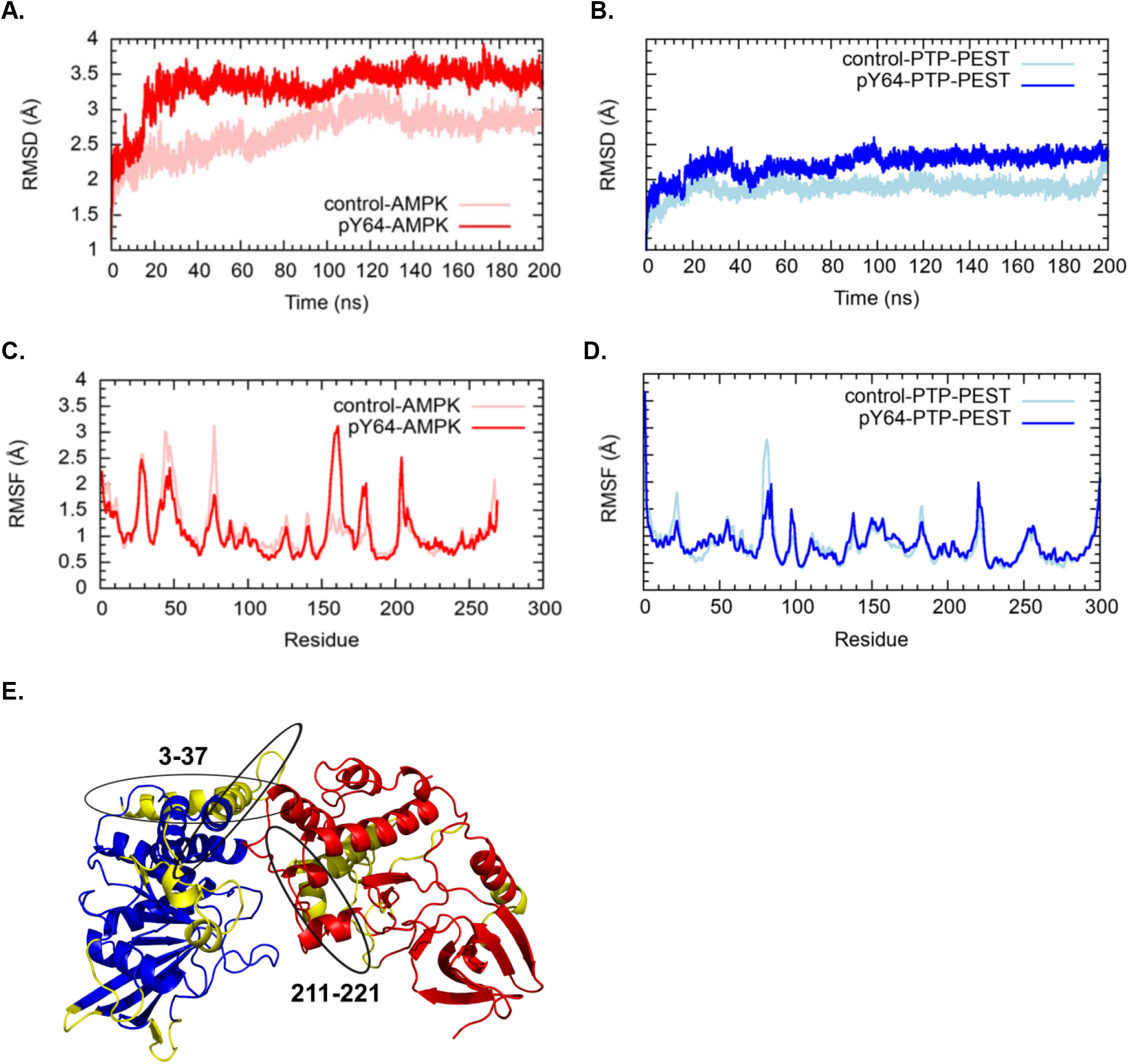
Molecular dynamic simulations for the PTP-PEST—AMPK(pY232) and PTP-PEST(pY64)—AMPK(pY232). (A) RMSD for AMPK from the the control (pink) and pY64 (red) complexes respectively. (B) RMSD for PTP-PEST from the control (light-blue) and pY64 (blue) complexes plotted for the length of the simulation. RMSF for (C) AMPK and (D) PTP-PEST in the control and pY64 complexes was plotted for the last 25ns of the simulation. (E) Ribbon representation of the simulated pY64 complex. The regions of PTP-PEST and AMPK showing changes in fluctuation between the control and pY64 complexes are highlighted in yellow.

**Figure 5.**
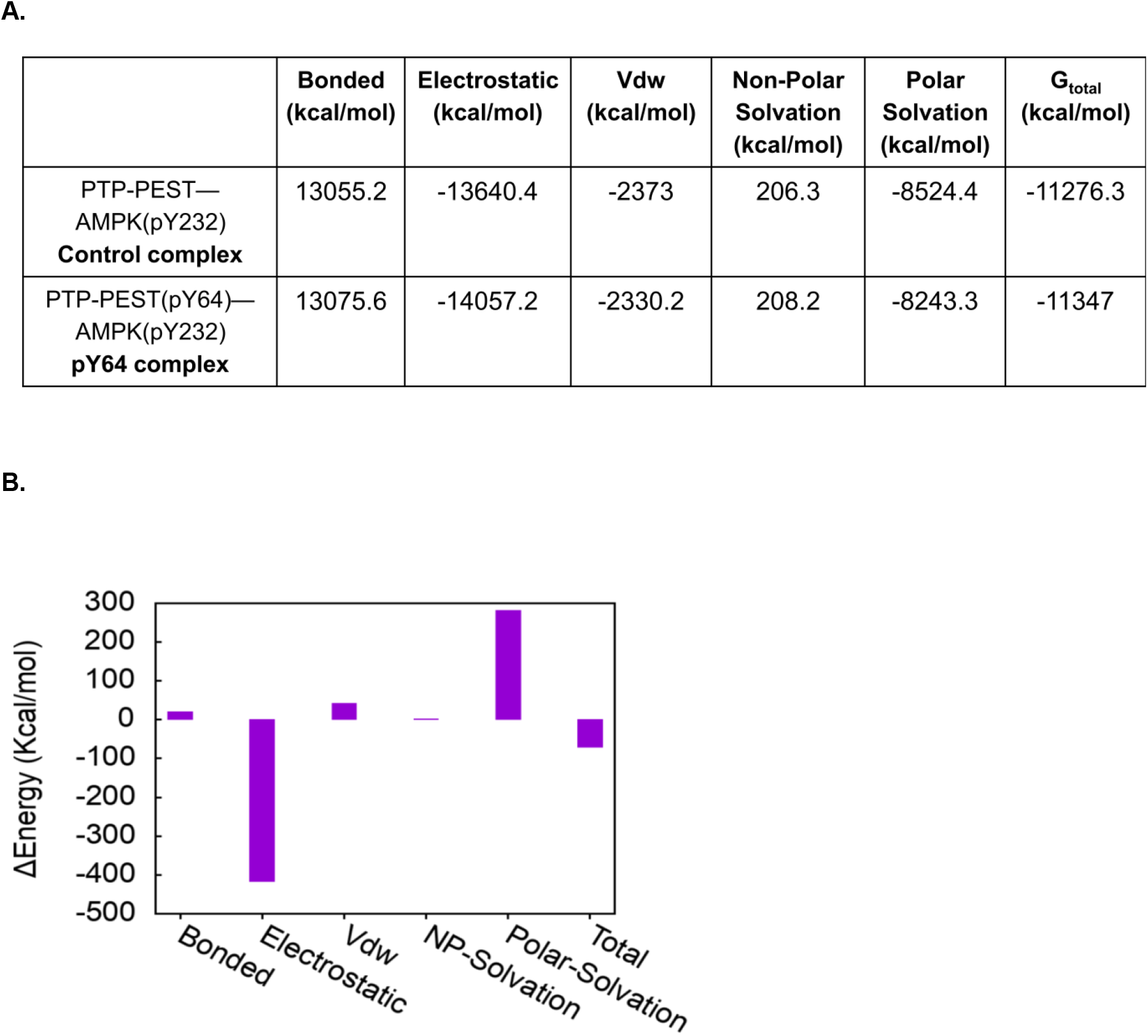
Free energy calculations of the simulated complexes. (A) Free energies, namely Bonded, Electrostatic, Van der Waals (VdW), Non-Polar and Polar Solvation as well as the total free energy (G_total_) of the control and pY64 complexes (B) Difference in free energies between the two complexes (ΔEpY64-ΔEcontrol) is plotted as ΔEnergy in Kcal/mol.

### 3.5 Phosphorylation of Y64 and rearrangement of local interaction networks

As with many protein-protein complexes, the interface of the PTP-PEST— AMPK(pY232) has many polar/charged residues and some are functionally relevant and have been implicated in substrate recognition as well^43^. The residues of relevance are the polar Y64, positively charged K142 of the β3-β4 loop and negatively charged D199 of the WPD loop as well as E137 of the β3-β4 loop in PTP-PEST (Supplementary Figure S1) and the negatively charged pY232 of AMPK. In the control complex, it is seen that K142 interacts with D199, located on the functionally relevant WPD loop of PTP-PEST, which in turn interacts with pY232 in AMPK at the complex interface (Figure 6, Supplementary Figure S9). The positively charged K142 forms two stable salt bridge interactions with negatively charged E137 and D199 in PTP-PEST (Figure 6A, C and D). Introduction of negative charges on Y64 after phosphorylation disrupts and reorganizes this electrostatic network. pY64 now interacts with K142, disrupting its interaction with E137 (Figure 6A, B and C). However, the ionic interaction between K142 and D199 is retained even after the phosphorylation of Y64 on PTP-PEST (Figure 6A, B and D). In addition to changes in the electrostatic interaction energies, introduction of the bulky phosphate group on Y64 also leads to steric rearrangements of the proximal residues. Both steric and electrostatic changes lead to the following scenario: a change in the conformation of K142 and the preservation of K142-D199 interaction (Figure 6A, D), which effectively draws the WPD loop towards the pY64 residue (Figure 6B, E, and Supplementary Figure S10) while the D199 residue continues to interact with the pY232 of AMPK (Figure 6F). A significant consequence of rearrangement of these residues (Figure 6G) is the reorientation of AMPK residue pY232 towards the complex interface upon phosphorylation of Y64 in PTP-PEST (Figure 6H and Supplementary Figure S11). The electrostatic interaction between two negatively charged pY232 and pY64 also shows a decrease in repulsive interaction energy between the two along the MD trajectory (Supplementary Figure S12C), and is consistent with the reorganization of the electrostatic network (Supplementary Figure S12). Thus, the electrostatic interactions of the two phosphosites with proximal positively charged residues are correlated with significant rearrangements occurring in the interactions in the pY64 system, where changes at the interface favour increased binding.

**Figure 6.**
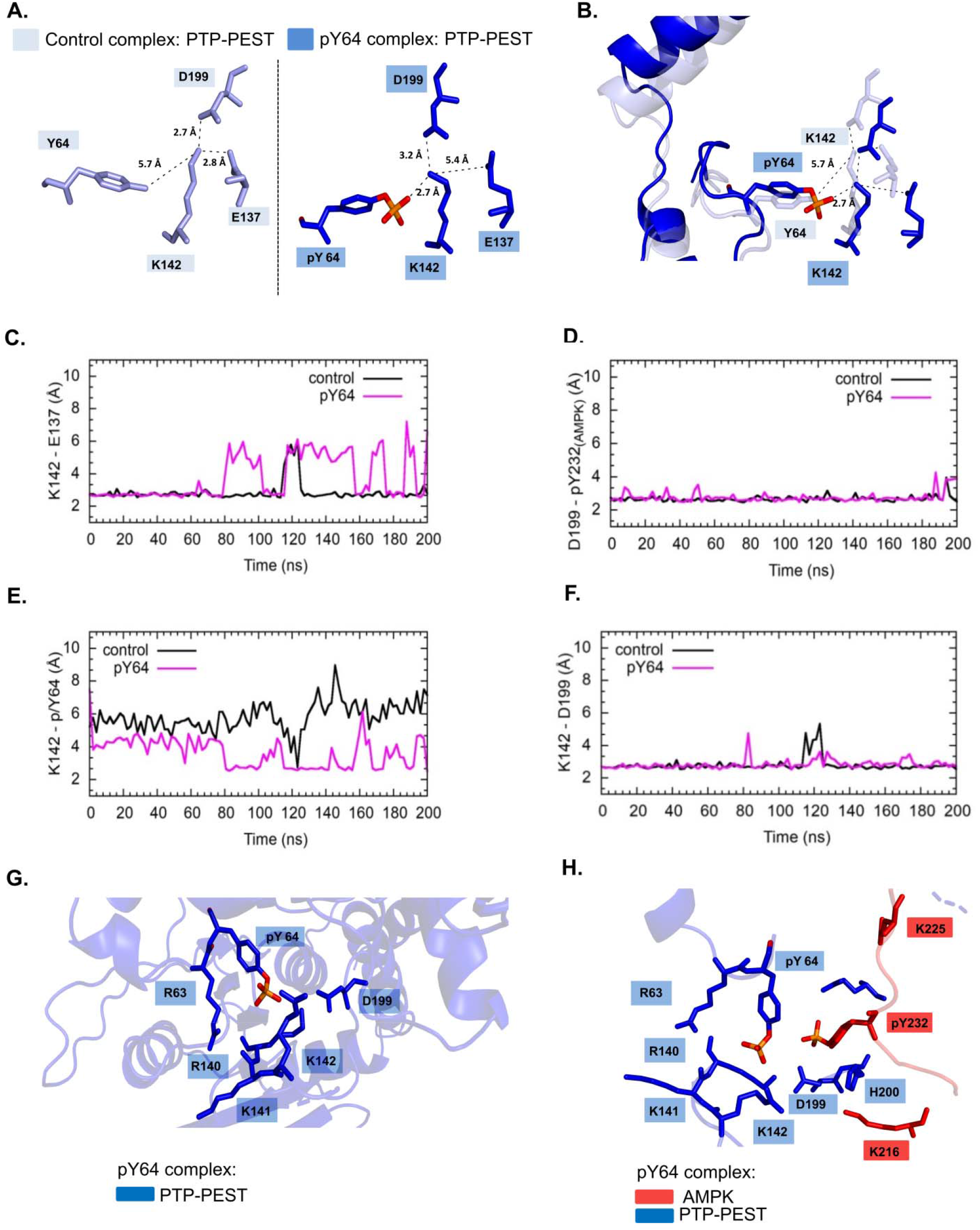
Local microenvironments of the control and pY64 complexes. Stick representation of the interactions of PTP-PEST residue K142 in the (A) control and pY64 complexes (B) Superposition of the interactions made by K142 in the two complexes. Distance plots between residues (C) E137 (PTP-PEST) - K142 (PTP-PEST), (D) D199 (PTP-PEST) - K142 (PTP-PEST), (E) K142 (PTP-PEST) - pY/Y64 (PTP-PEST) and (F) pY232 (AMPK) - D199 (PTP-PEST), where the black line indicates the control complex and the pink line indicates the pY64 complex (G) Stick representation of the residues in the vicinity of pY64 on PTP-PEST (blue) and (H) pY232 of AMPK (red) in the pY64 complex.

The residue H200 in PTP-PEST (on WPD loop) is reported to play an important role in substrate selectivity and our simulations suggest that in the pY64 complex, the conformation of H200 changes significantly (Supplementary figure S13A and B). A new interaction with K225 (AMPK), in the form of a hydrogen bond interaction emerges, further contributing to the increased binding (Figure 7A and Figure 7B). Comparison of the control and pY64 simulations show that the conformation of H200 residue is locked throughout the simulation time scale in the pY64 complex, unlike the control system where a significant reorientation of H200 is observed (Figure 7B, C and Supplementary Figure S13). The reorganization of the interaction can also be quantified by Protein network analysis (Figure 8A, B). The data show that in the pY64 complex, the closely interacting community definitions change and a new community spanning relevant residues at the complex interface emerges. Taken together, all these results represent the atomic level changes in conformation and interactions that manifest as increased binding of the complex upon phosphorylation of Y64 of PTP-PEST.

**Figure 7.**
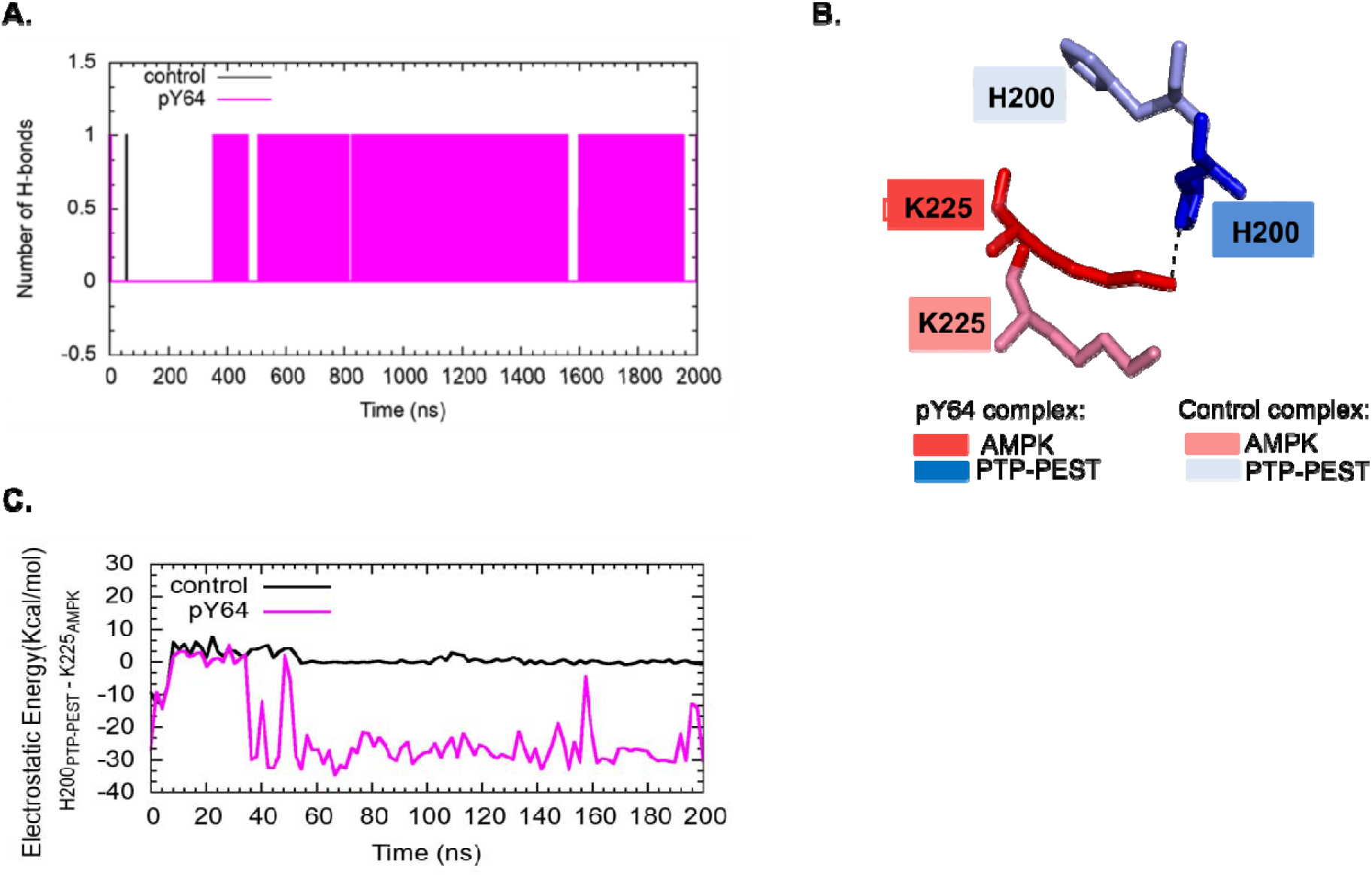
Hydrogen bond dynamics of H200 upon the phosphorylation of Y64 on PTP-PEST. (A)Comparison of the change in H-bonds formation by H200 across 200ns of simulation upon phosphorylation of pY64 on PTP-PEST, here the black line indicates bonds formed by H200 in the control complex and the pink line indicates bond formation by H200 in the pY64 complex. (B) Hydrogen bond formation between side chains of H200 and K225. Light blue and pink denote PTP- PEST and AMPK respectively, from the control complex, whereas blue and red denote PTP-PEST and AMPK respectively, from the pY64 complex. (C)Electrostatic energy graph for residues K225 (AMPK) and H200 (PTP-PEST) in the control (black) and pY64 4 (pink) complexes.

**Figure 8.**
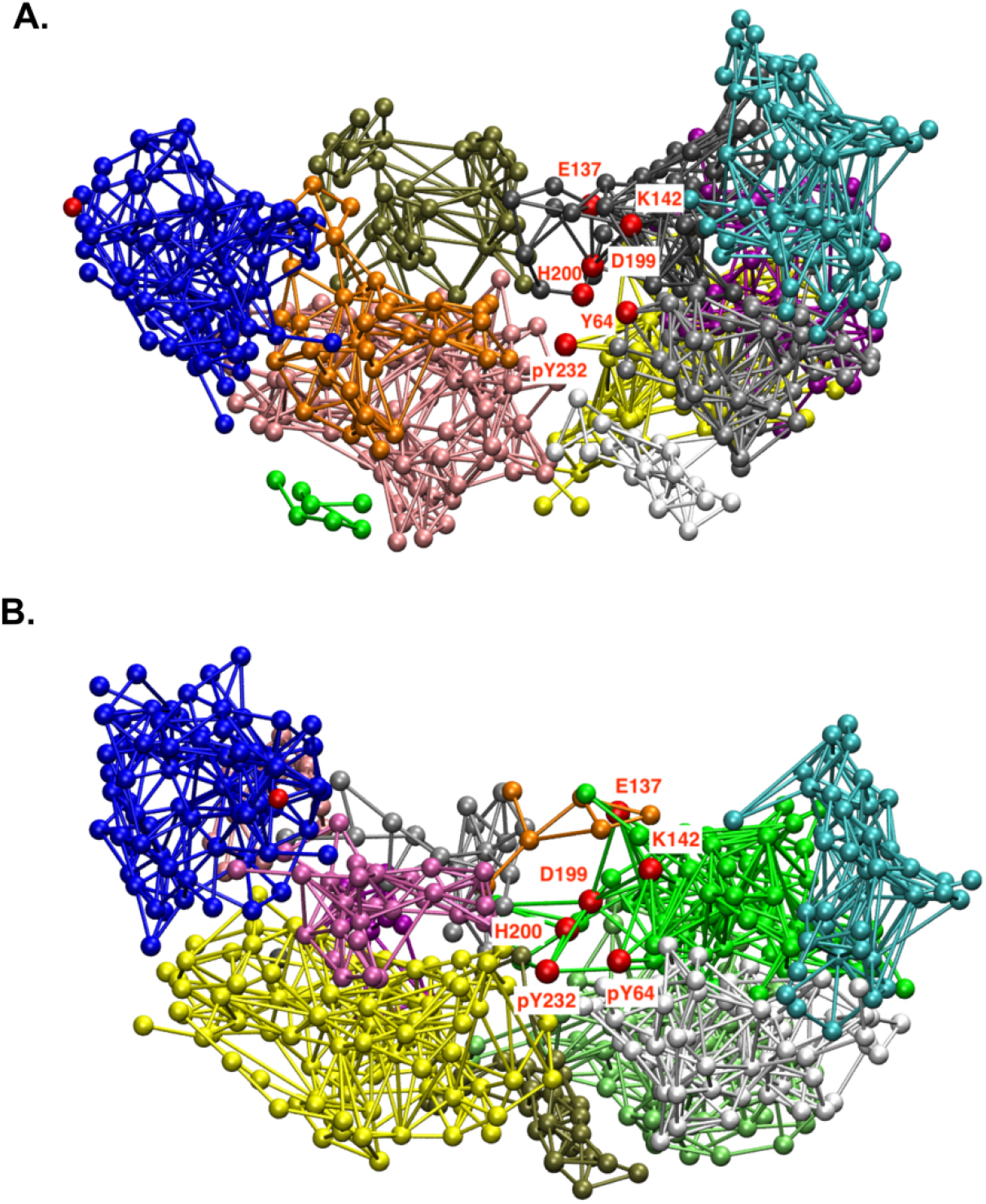
Differences in the interaction networks of the control and pY64 complexes. Protein network analysis of (A) control PTP-PEST—AMPK(pY232) complex and (B) pY64 PTP-PEST(pY232)—AMPK(pY64) complex with the residues of interest highlighted in red

## 4 Discussion

In this study we used computational analyses to identify a potential tyrosine dephosphorylation site on AMPK that is targeted by PTP-PEST. Docking studies to generate complexes between AMPK and PTP-PEST, followed by analyses of binding energy, interface area and non-bonded interactions, were used to validate the choice of residue Y232 of AMPK as the most likely phosphosite. The phosphotyrosine residues in complexes of AMPK phosphorylated at the other potential sites, namely, Y104 and Y283, were not positioned near the catalytic P-loop, as would be required in an enzyme-substrate complex. Although the overall geometry of the interactions in the PTP-PEST—AMPK(pY232) complex was as expected for a productive complex, the rigid docking protocol used here does not account for any local conformational changes that may occur *in vivo*. Furthermore, examination of the active site shows that H274, a crucial residue involved in determining the substrate specificity of PTP-PEST is present at the interface of the complex. An earlier study proposed that H274 in PTP-PEST forms the ‘pY+1’ pocket and interacts with residues C-terminal to the substrate phosphotyrosine^43^. As expected, the residue at this position is significantly divergent across homologous PTPNs. For instance, human PTPN18, PTPN2, PTPN6, and PTPN7, contain proline, methionine, serine and glycine, respectively, at this position^8^. In our model complex, H274 is indeed located at a distance of 3.7 Å from the C-terminal residue E235 of AMPK (Supplementary Figure S14). Together, these observations strongly support the notion that our proposed model most likely represents a valid complex. Atomistic molecular dynamics simulations were performed to check the stability of the PTP-PEST—AMPK(pY232) complex and to understand the interaction network at the complex interface. The changes in RMSD and RGyr indicated a stable complex at 100ns of simulation. It is also known that the enzymatic activity of PTP-PEST is regulated by phosphorylation^8^. Interestingly, Y64, a key catalytic residue of PTP-PEST is a known phosphosite and is located at the interface of the complex. Mutation of this residue displayed markedly lowers activity compared to the wild type^21^. Considering that the residue pY232 of AMPK is located in the immediate neighbourhood of Y64 of PTP-PEST, it was expected that the introduction of negative charges via phosphorylation of Y64 can potentially affect the electrostatic network interactions at the interface and therefore, the energetics of the complex binding. Thus, additional simulations including free energy calculations were performed to probe its effect on the overall structure and stability of the PTP-PEST-AMPK(pY232) complex.

The MD simulations suggest that the phosphorylated PTP-PEST(pY64) binds more strongly with AMPK(pY232) compared to the unphosphorylated PTP-PEST system. The increased binding primarily originates from altered interactions arising out of the rearrangement of charged residues in PTP-PEST located at the interface of the complex. The functionally relevant WPD loop (residues 197-199) on PTP-PEST is highly conserved in classical PTPs. Residue D199 on this loop is a known proton donor in the catalysis and is located proximal to the complex interface. In the control unphosphorylated complex, D199 is seen to interact with both pY232 of AMPK and K142 of PTP-PEST. The phosphorylation of Y64 of PTP-PEST induces a change in the electrostatic network around this residue. The negatively charged pY64 now forms a stable interaction with the positively charged K142 which continues to interact with D199 even after phosphorylation of Y64. This change in the interaction network results in the movement of the WPD loop towards the pY64 residue on PTP-PEST. Simultaneously, the pY232 residue of AMPK moves closer towards the interface. The resulting increase in favourable electrostatic interaction energy between AMPK and PTP-PEST possibly contributes to increased binding affinity of the complex.

In addition, a new interaction between H200 of PTP-PEST and K225 of AMPK was observed upon phosphorylation of Y64 on PTP-PEST further contributing to the increased binding in the complex. H200 is crucial for accurate positioning of the proton donor during catalysis and the mutation of this residue is known to significantly impair dephosphorylation of phosphorylated substrates^8^. Notably, H200 participates in charge-charge interactions with the substrate phosphopeptides, as was also observed in the case of the PTP-PEST—AMPK (pY232) complex. Although the catalytic cleft is largely conserved in PTPNs, the residue H200 is specific to PTP-PEST and is not found in homologous phosphatases such as PTPN18 (substituted by N/R in PTPN18, Supplementary Figure S6). The movements of the WPD loop observed during our simulation suggest that potential conformational changes occur in this loop after phosphorylation of Y64 of PTP-PEST, affecting its catalytic activity, in that movement of the WPD loop of PTP-PEST and pY232 of AMPK further stabilizes this complex. Whether such an increased binding results in altered enzymatic kinetics or can potentially lead to substrate trapping is an interesting future direction to investigate. Our earlier experimental studies have shown that AMPKα interacts with PTP-PEST under normoxic condition and that this interaction is impaired in hypoxia-treated cells^9^. The observations from the current simulations indicate an increased binding between AMPKα and PTP-PEST upon phosphorylation of Y64 on PTP-PEST. Given that protein phosphorylation is a ubiquitous mechanism for the activation and signalling in cellular processes, it is tempting to speculate that the regulation of AMPK by PTP-PEST under physiological conditions is mediated by the phosphorylation/dephosphorylation states of both Y232 in the AMPKα subunit and of residue Y64 in the catalytic domain of PTP-PEST. These hypotheses remain to be validated by *in vitro* and *in vivo* measurements.

## Supporting information

All Supplementary files

## Acknowledgements

Amrutha Manikandan would like to acknowledge the Ministry of Human Resources and Development (MHRD) for her fellowship. The simulations were carried out on the supercomputing machines Annapurna and Nandadevi at The Institute of Mathematical Sciences.

## Data Availability Statement

The data would be made available on request from the authors.

