## Supplementary material for "*In-silico* identification of Tyr232 in AMPKα2 as a dephosphorylation site for the protein tyrosine phosphatase PTP-PEST": All Supplementary files

**Running title:** Interaction of PTP-PEST with AMPKα2

*** Corresponding Authors**

Madhulika Dixit, Ph.D.

Department of Biotechnology, Bhupat and Jyoti Mehta School of Biosciences

Indian Institute of Technology Madras

Narayanan Manoj, Ph.D.

Department of Biotechnology, Bhupat and Jyoti Mehta School of Biosciences

Indian Institute of Technology Madras

**Supplementary Figures**

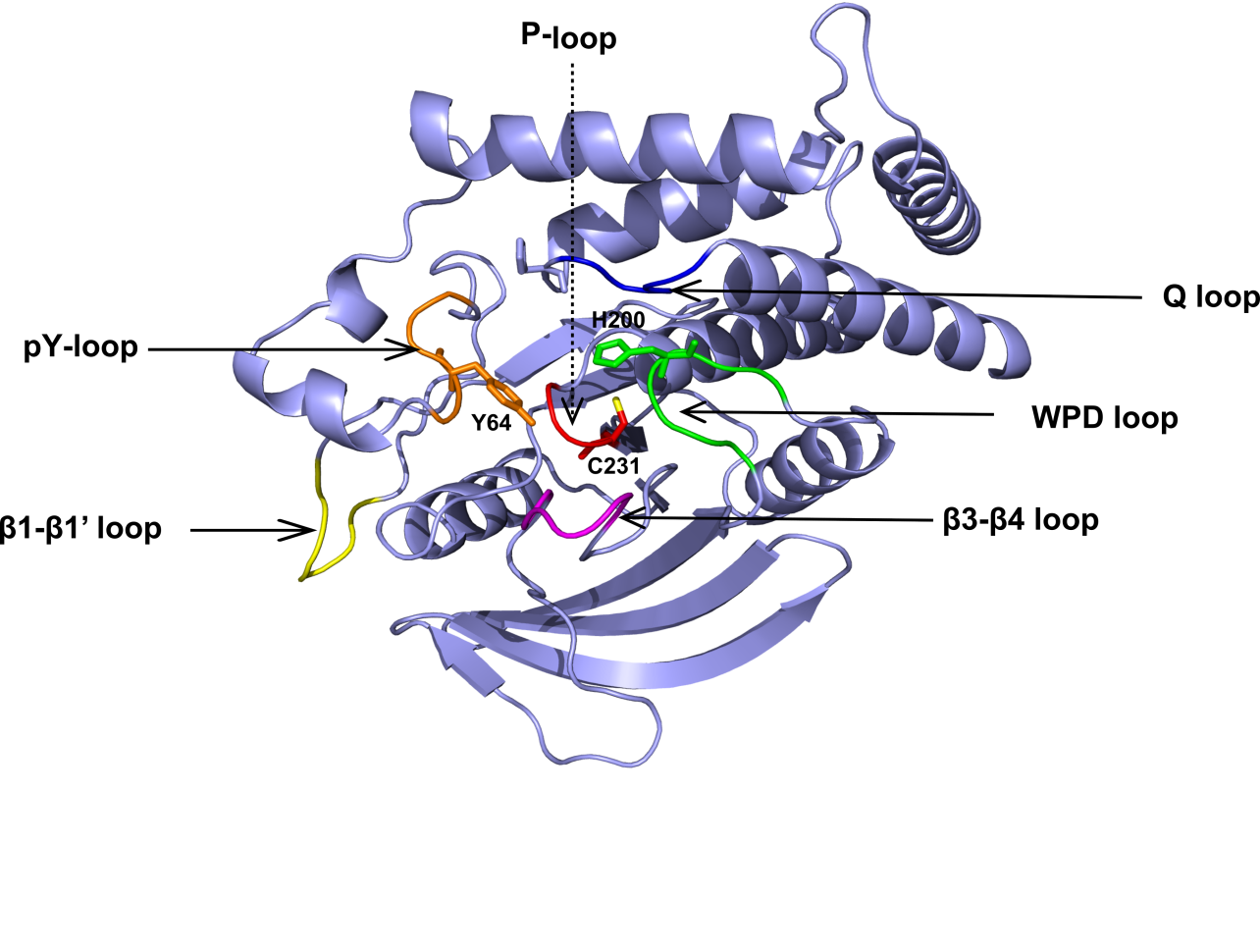

**Supplementary Figure S1**

**Supplementary Figure S1. Crystal structure of the human PTP-PEST catalytic domain.** Ribbon diagram of the catalytic domain of PTP-PEST with the active site flexible loops highlighted in orange (pY), yellow (β1-β1’), dark blue (Q), green (WPD), pink (β3-β4) and red (P) (PDB ID: 5HDE). Y64 of the pY loop and the catalytic cysteine C231 are shown in stick representation.

**Supplementary Figure S2**

**A.**

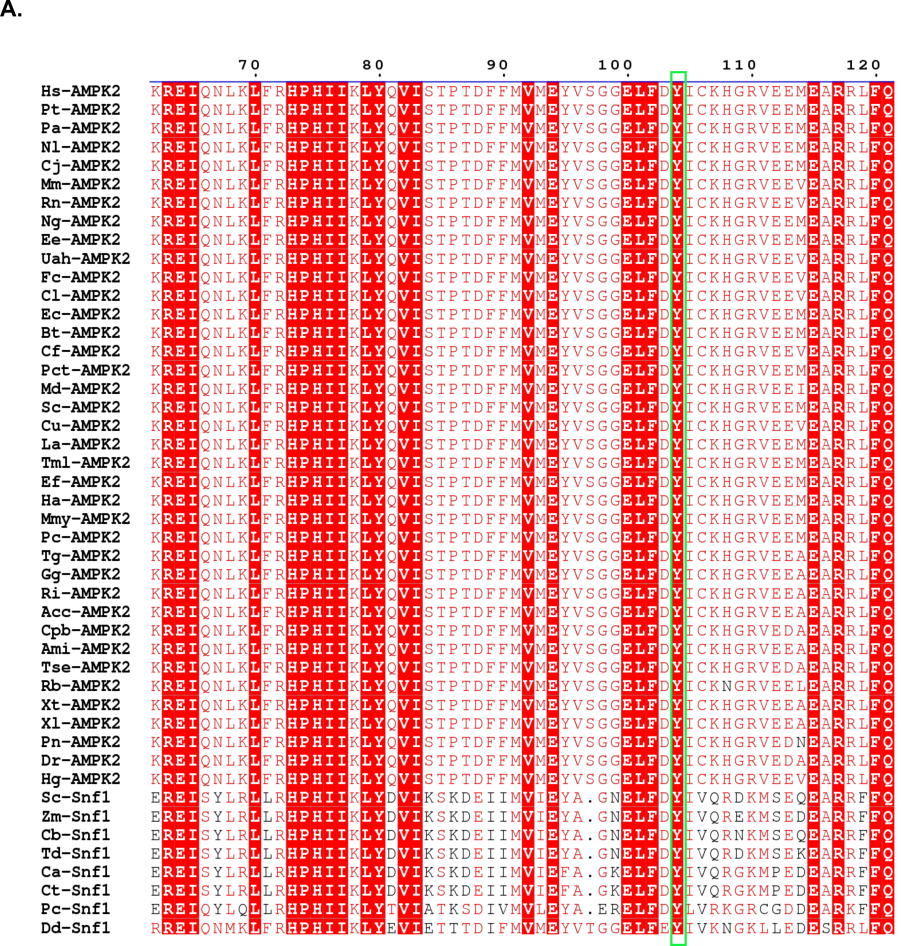

**B.**

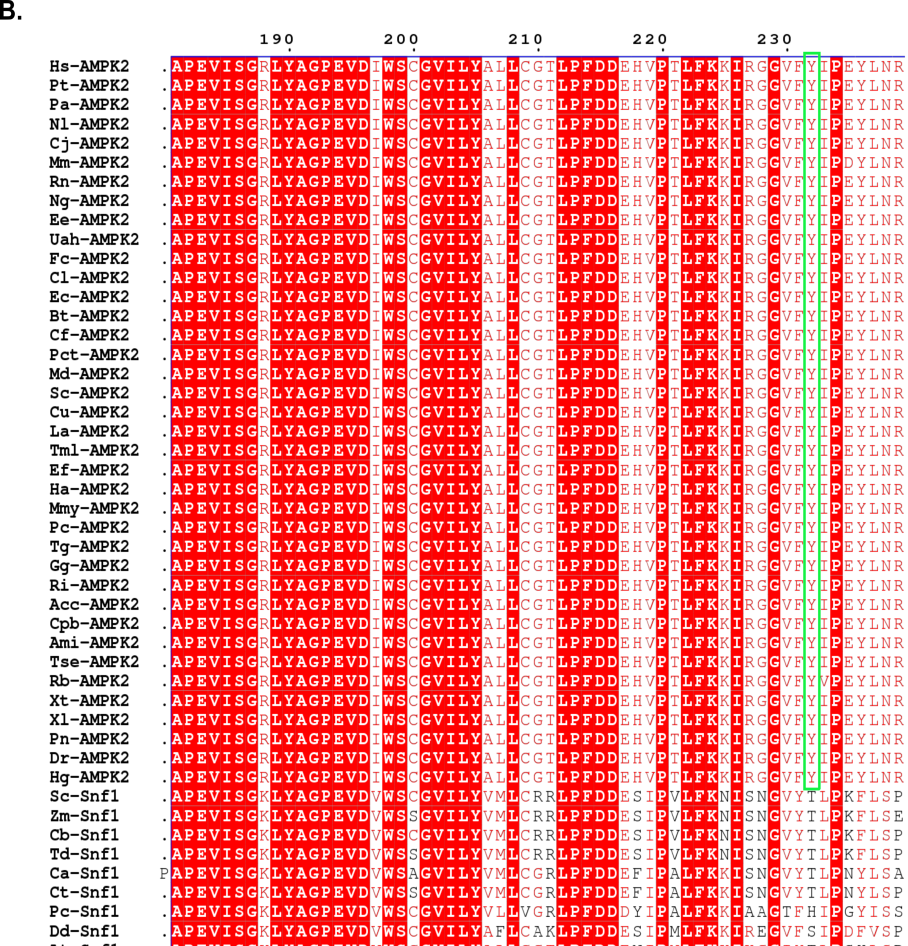

**C.**

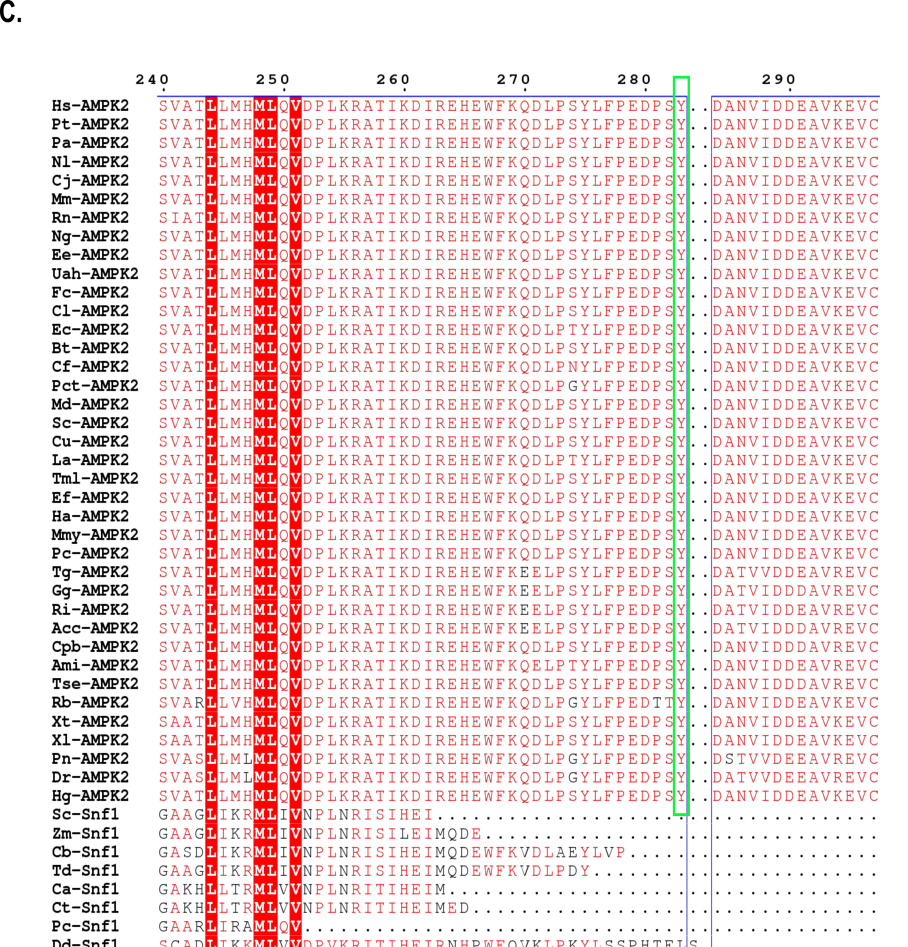

**Supplementary Figure S2. Multiple Sequence Alignment of AMPKα2 with a red arrow marking the start of invertebrate organisms**. (A) Sequence conservation of Y104 across organisms is marked in green. (B) Sequence conservation of Y232 across vertebrates is marked in green (C) Sequence conservation of Y283 across vertebrates is marked in green

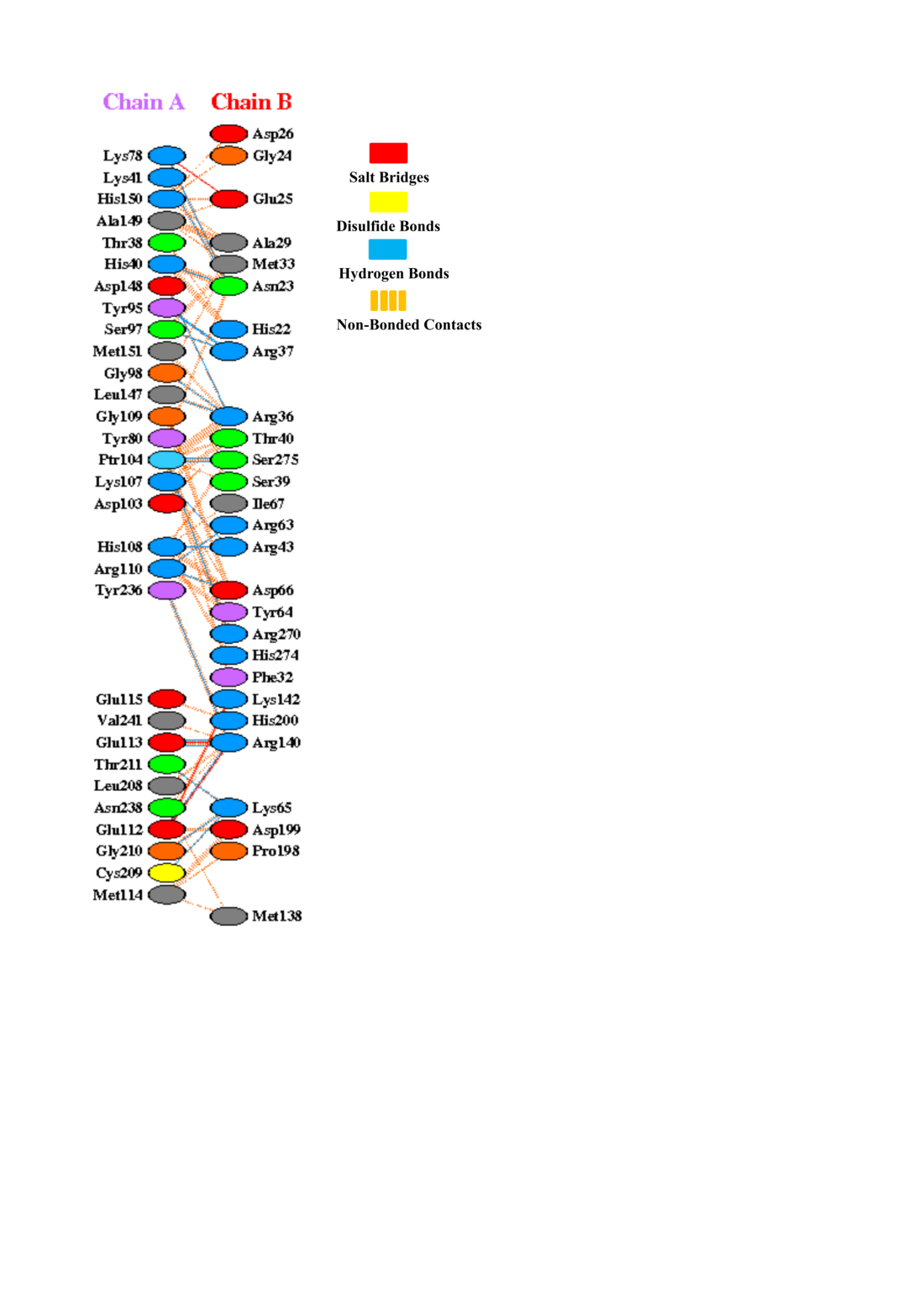

**Supplementary Figure S3**

**Supplementary Figure S3. PDBsum illustration of the protein-protein interface interactions of the PTP-PEST— AMPK(pY104) complex.** Here chain A refers to AMPK and chain B refers to PTP-PEST. The red dotted lines refer to salt bridges, yellow lines refer to disulfide bonds, blue lines refer to hydrogen bonds and the orange dashes refer to other non-bonded contacts.

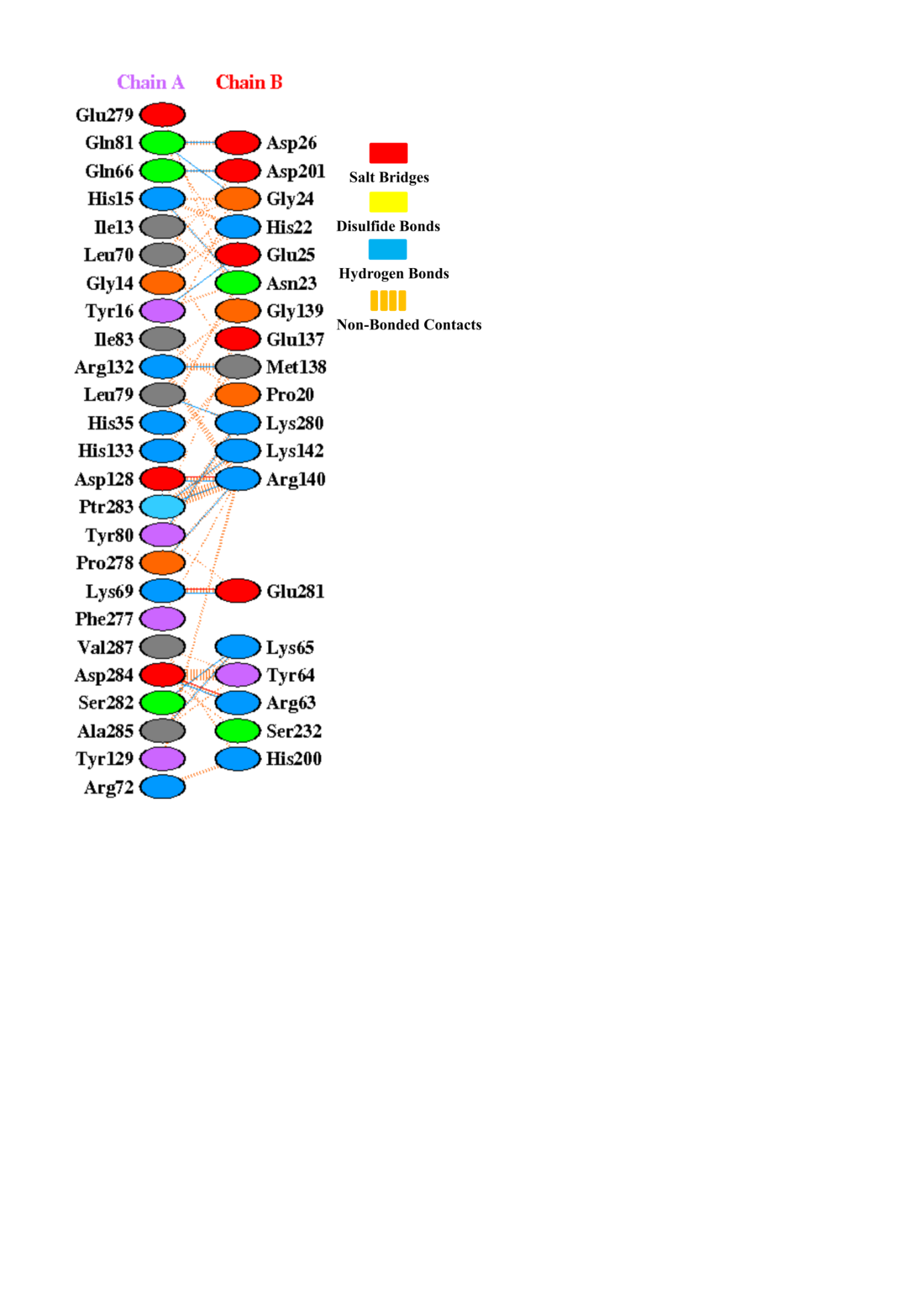

**Supplementary Figure S4**

**Supplementary Figure S4. PDBsum illustration of the protein-protein interface interactions of the PTP-PEST—AMPK(pY283) complex.** Here chain A refers to AMPK and chain B refers to PTP-PEST. The red dotted lines refer to salt bridges, yellow lines refer to disulfide bonds, blue lines refer to hydrogen bonds and the orange dashes refer to other non-bonded contacts.

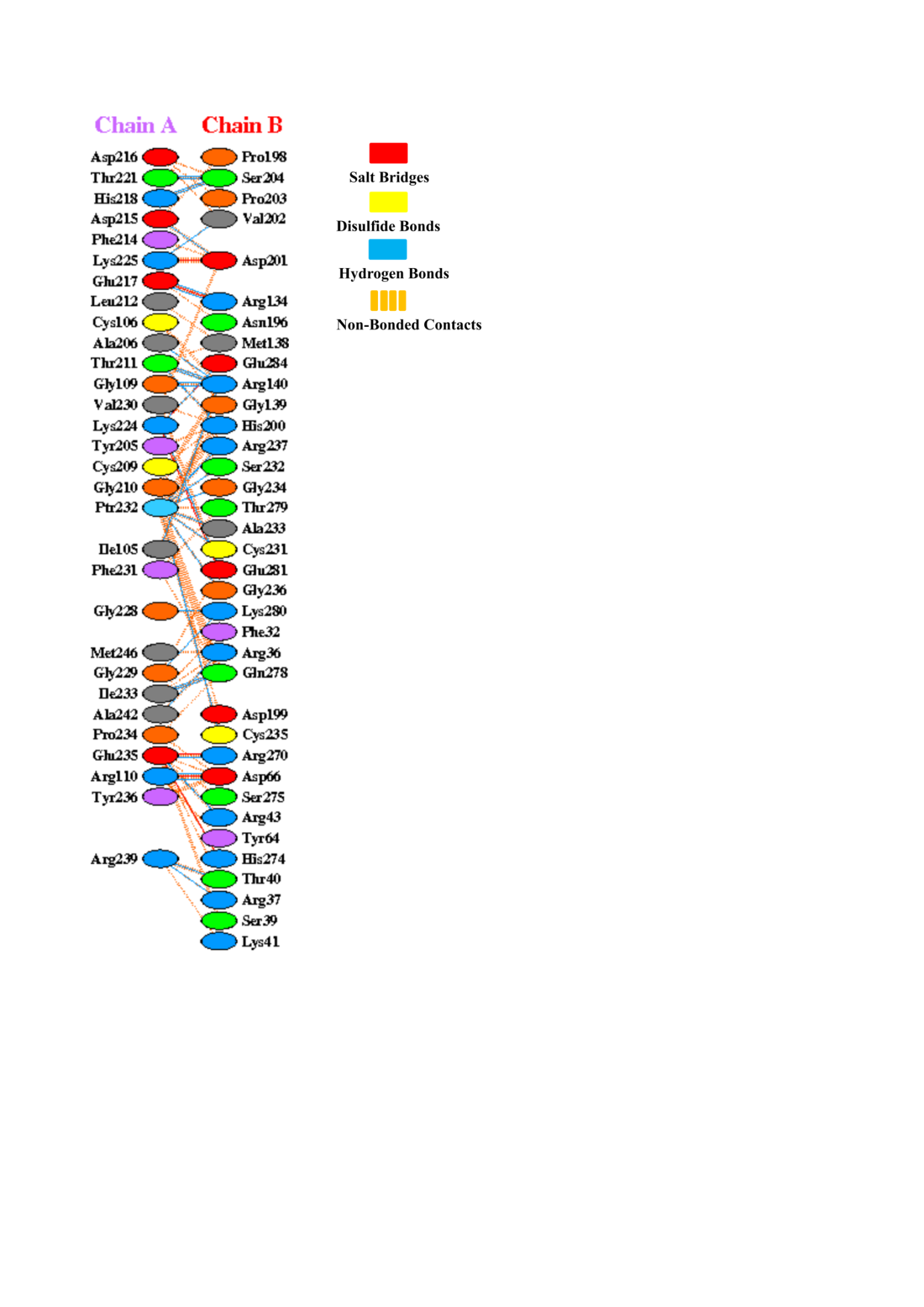

**Supplementary Figure S5**

**Supplementary Figure S5. PDBsum illustration of the protein-protein interface interactions of the PTP-PEST—AMPK(pY232) complex.** Here chain A refers to AMPK and chain B refers to PTP-PEST. The red dotted lines refer to salt bridges, yellow lines refer to disulfide bonds, blue lines refer to hydrogen bonds and the orange dashes refer to other non-bonded contacts.

**Supplementary Figure S6**

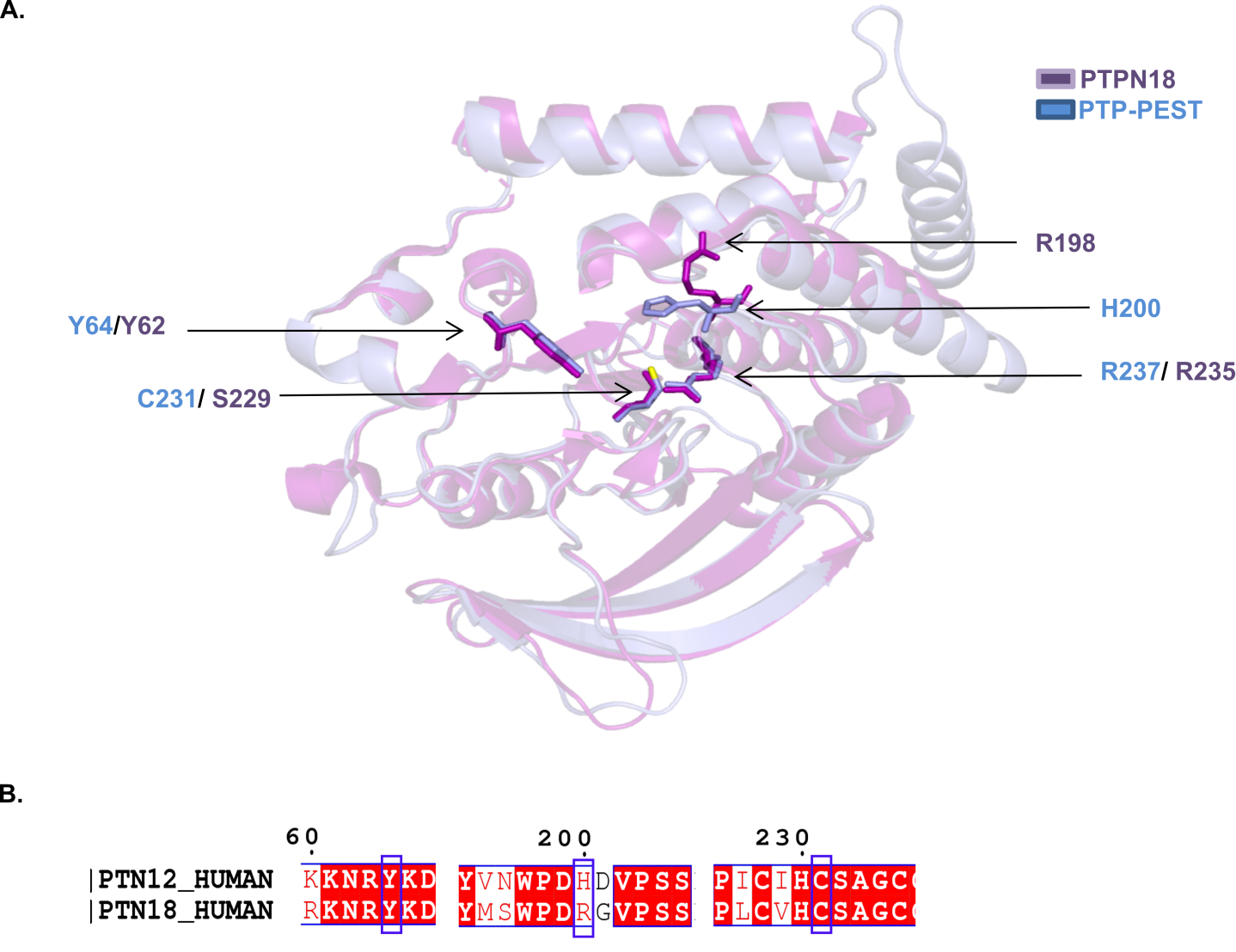
**A.**

**B.**

**Supplementary Figure S6. Structural similarities between PTP-PEST and PTPN18.** (A) Ribbon representation of the superposition of PTP-PEST (light-blue) with PTPN18 (purple). The catalytic cysteine C231, H200, Y64 as well as the catalytic arginines R237 of PTP-PEST and residues corresponding to these positions on PTPN18, are in stick representation (B) Sequence alignment between PTP-PEST and PTPN18, with blue boxes highlighting the conservation of PTP-PEST residues, namely Y64 and C231 as well as the lack of H200 in PTPN18.

**Supplementary Figure S7**

**A. B.**

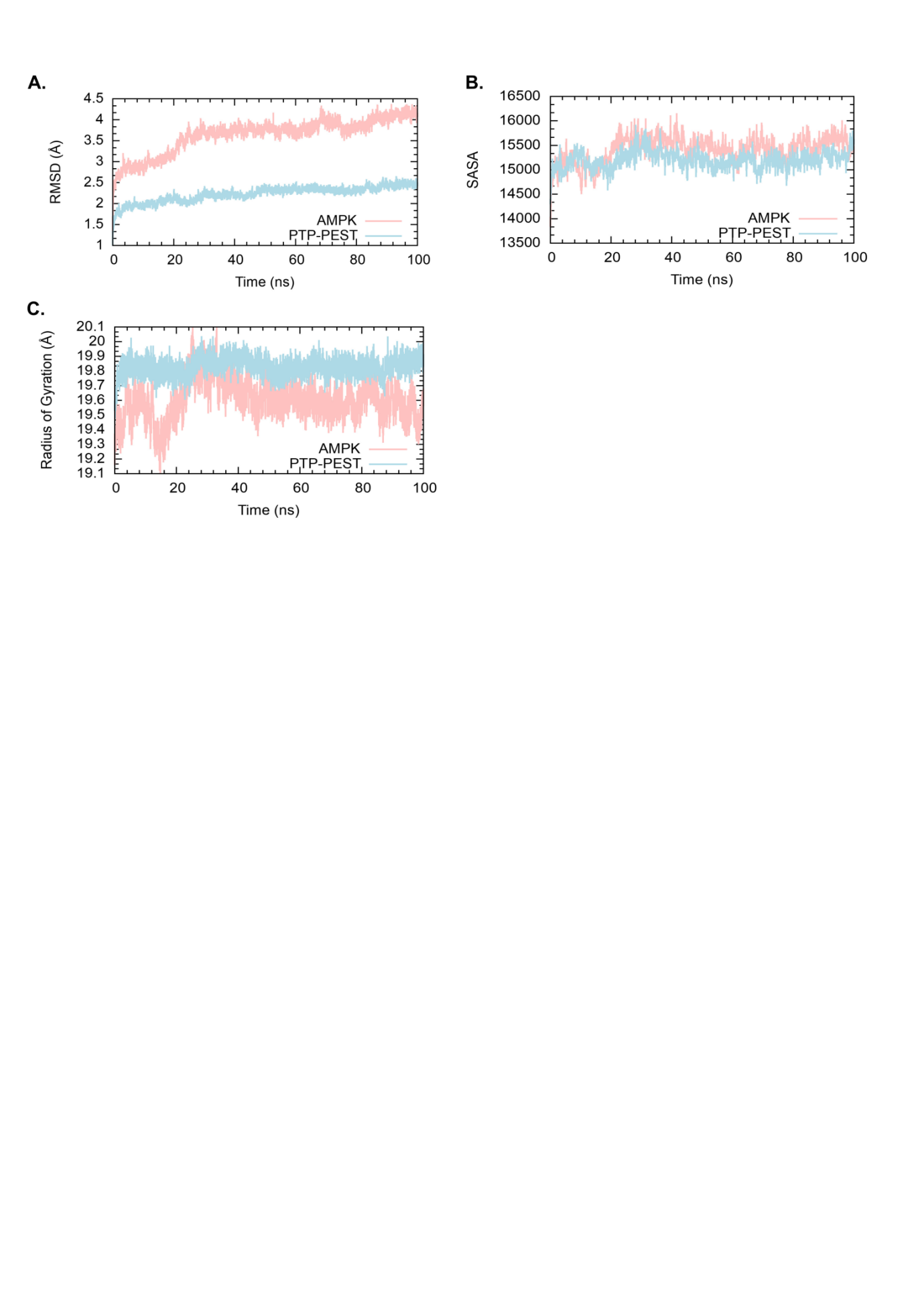
**C.**

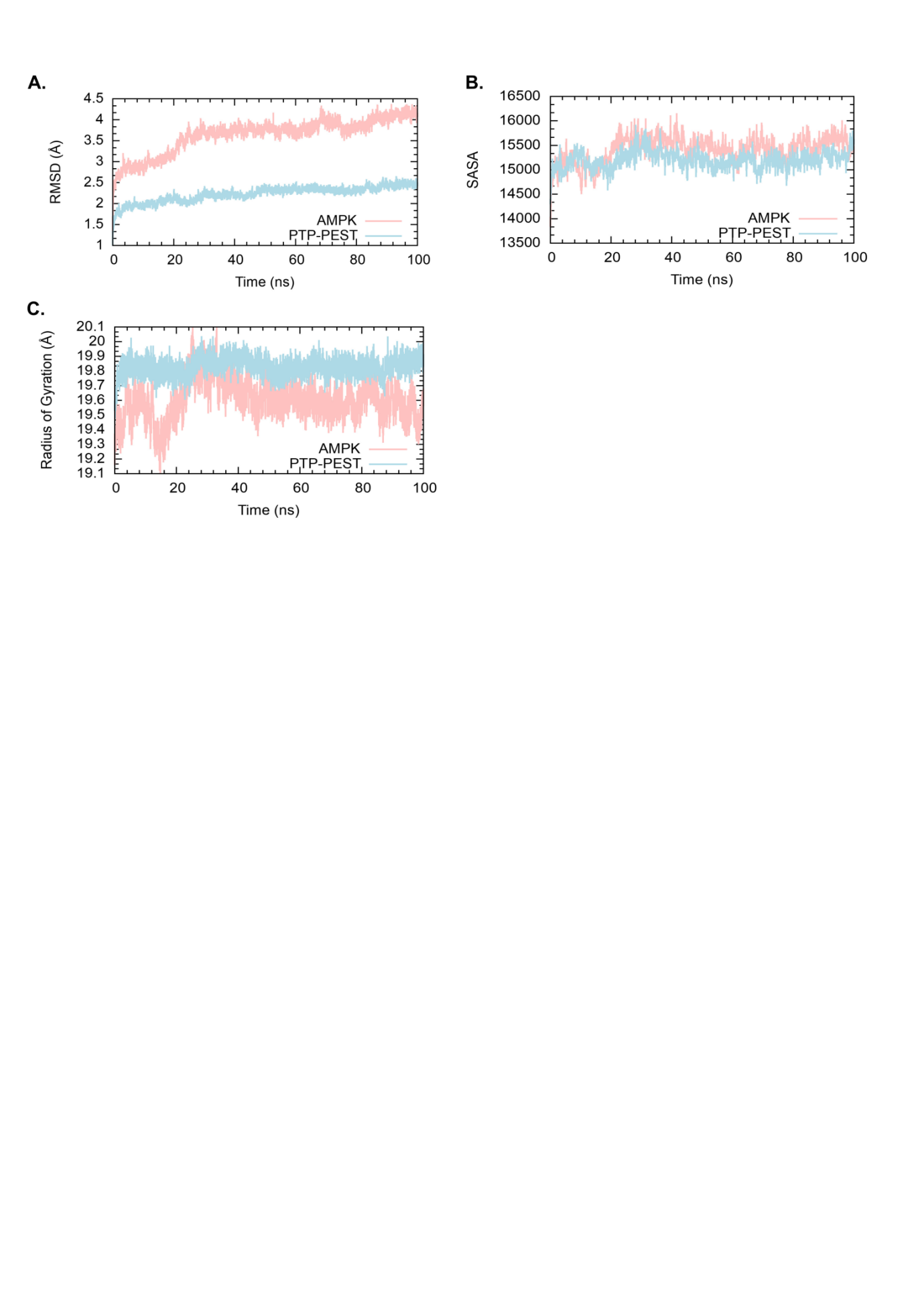

**Supplementary Figure S7.** **Trajectory Analysis of the simulated PTP-PEST —AMPK (pY232) complex.** (A) RMSD (B) SASA and (C) Rg analyses were done to check for the stability of the docked PTP-PEST-AMPK (pY232) complex after 100ns of simulation.

**A. B.**

**Supplementary Figure S8**

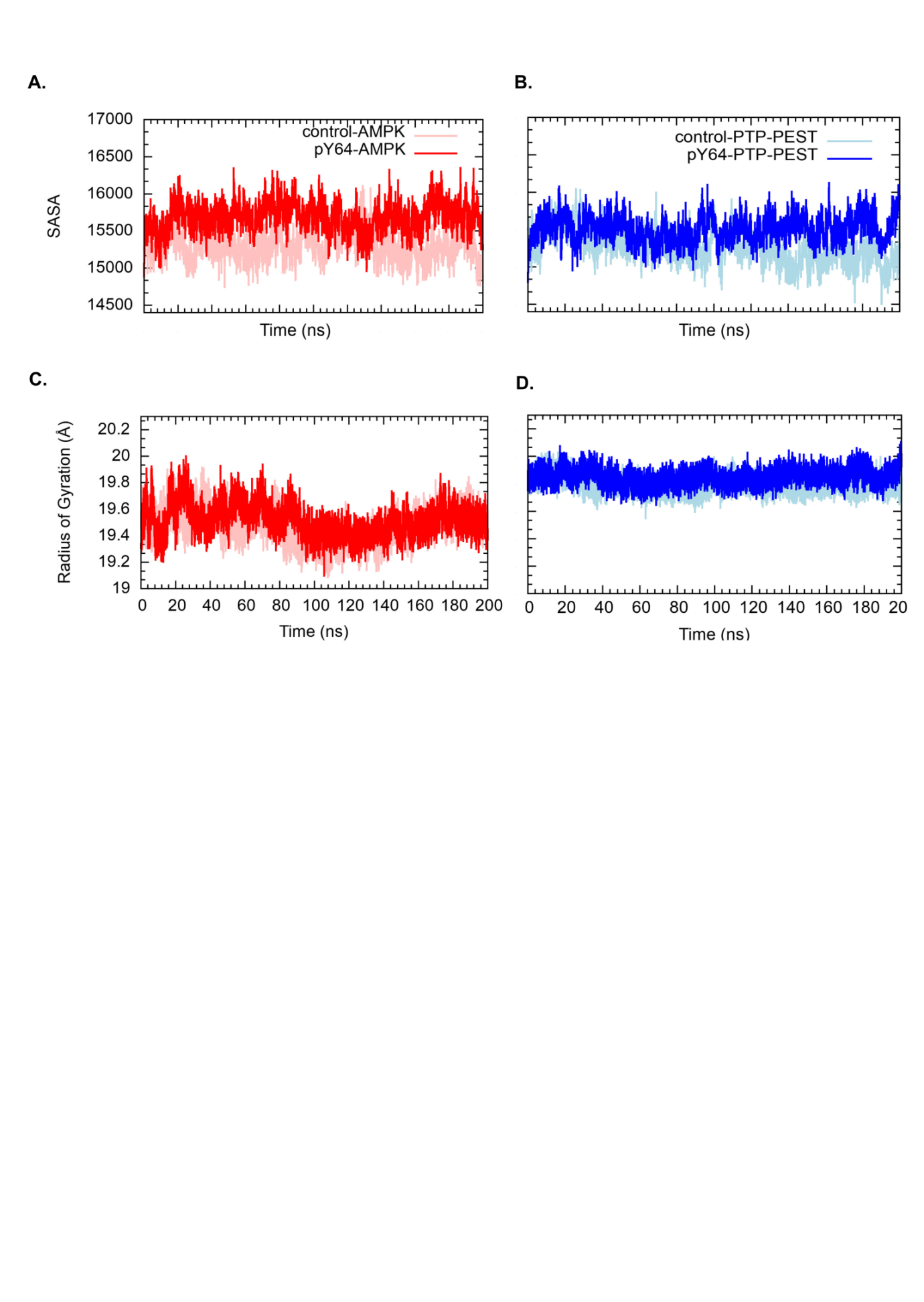

**C. D.**

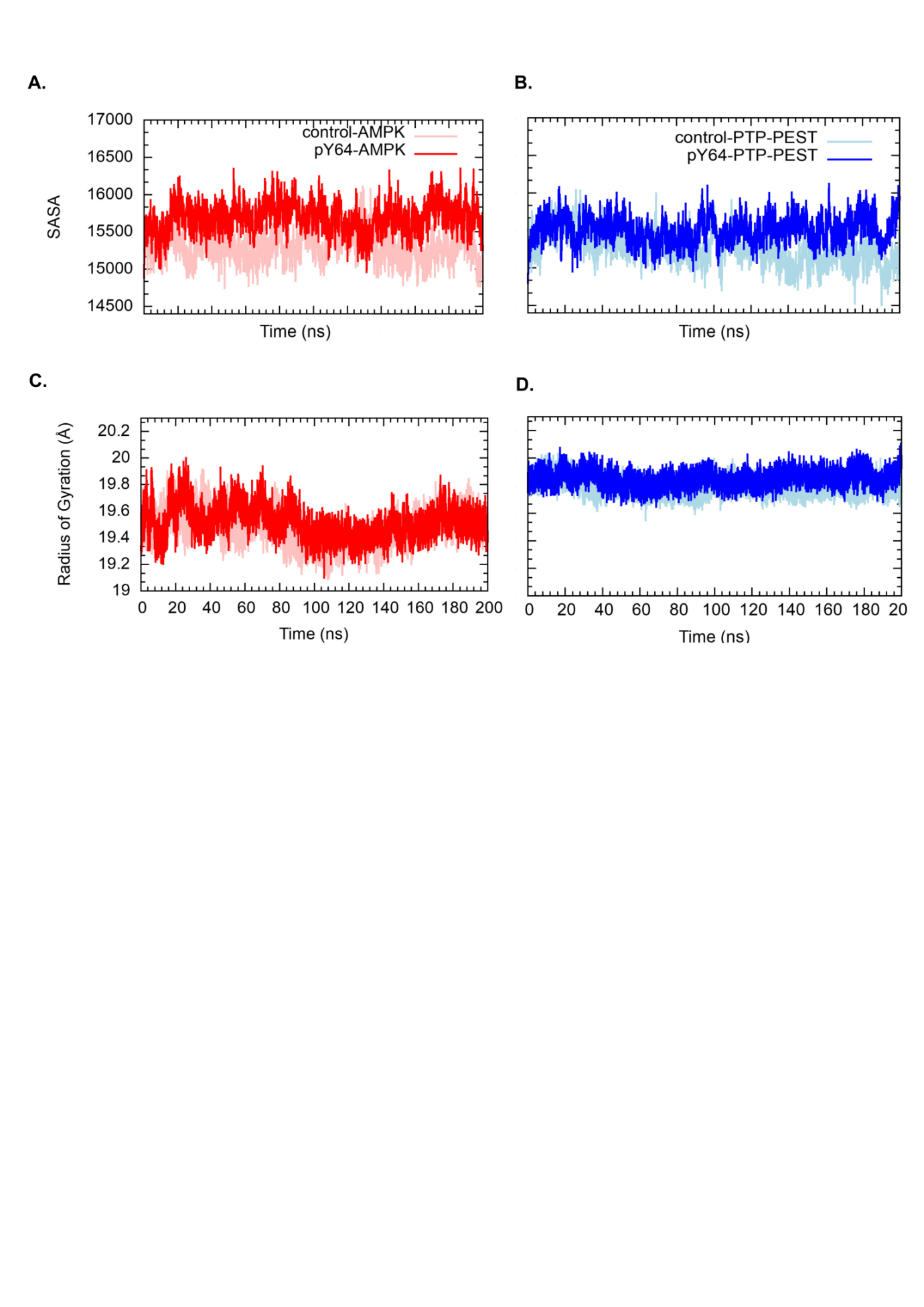

**Supplementary Figure S8.** **Evaluation of the control complex: PTP-PEST—AMPK(pY232) vs the pY64 complex: PTP-PEST(pY64) —AMPK(pY232).** Solvent Accessible Surface Area (SASA) plotted for (A) AMPK in the control (pink) and pY64 (red) complexes and SASA for (B) PTP-PEST in the control (light blue) and pY64 (blue) complexes. Radius of gyration (Rg) for (C) AMPK in the control (pink) and pY64 (red) complexes and Rg for (D) PTP-PEST in the control (light blue) and pY64 (blue) complexes plotted over the length of the simulation.

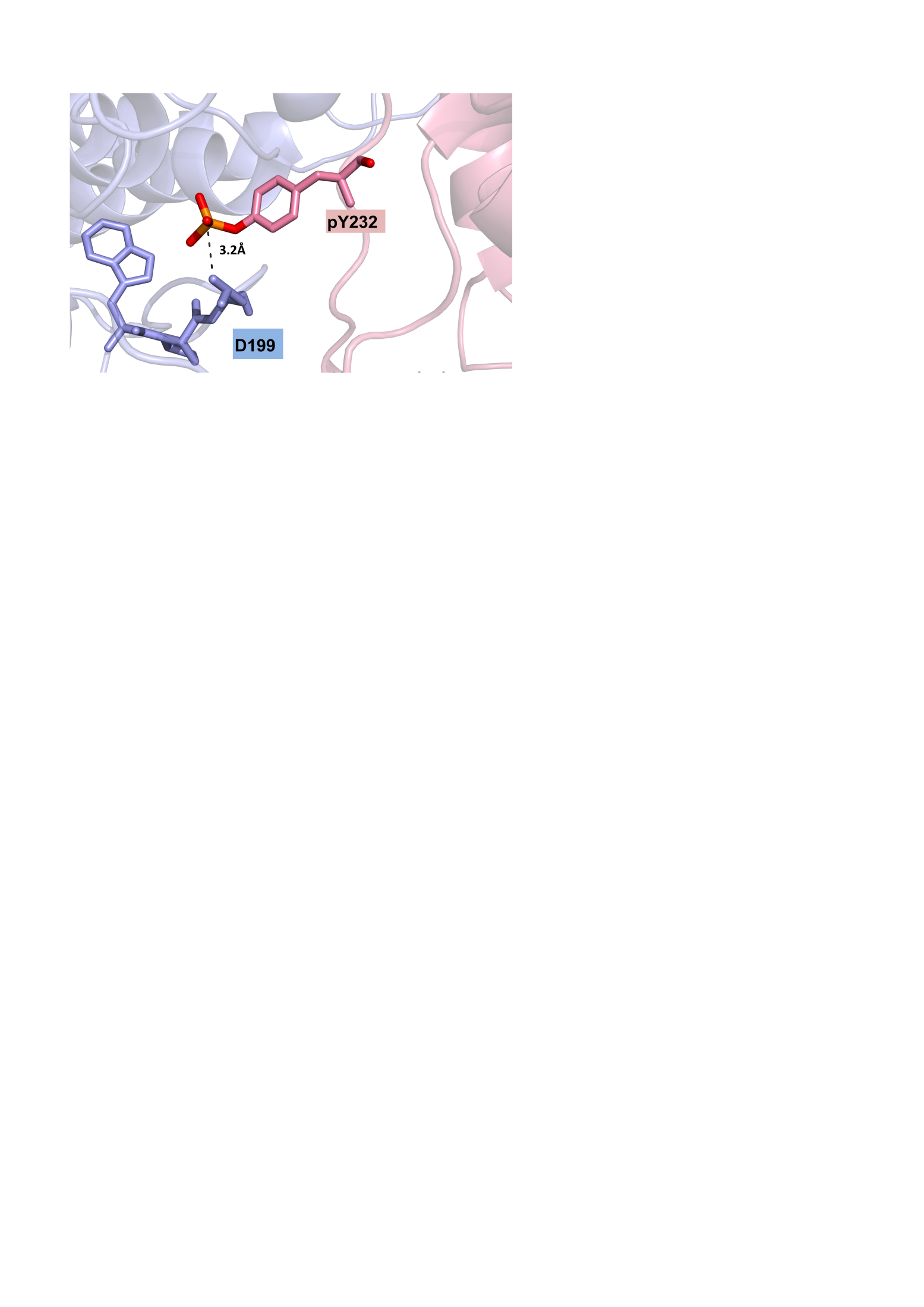

**Supplementary Figure S9**

**Supplementary Figure S9**. **Interface of the control PTP-PEST —AMPK(pY232) complex.** The interface between PTP-PEST (light blue) and AMPK (pink) shows the interaction between D199 (stick representation) on the WPD loop of PTP-PEST and pY232 (stick representation) on AMPK, that is retained after 190ns of simulation.

**Supplementary Figure S10**

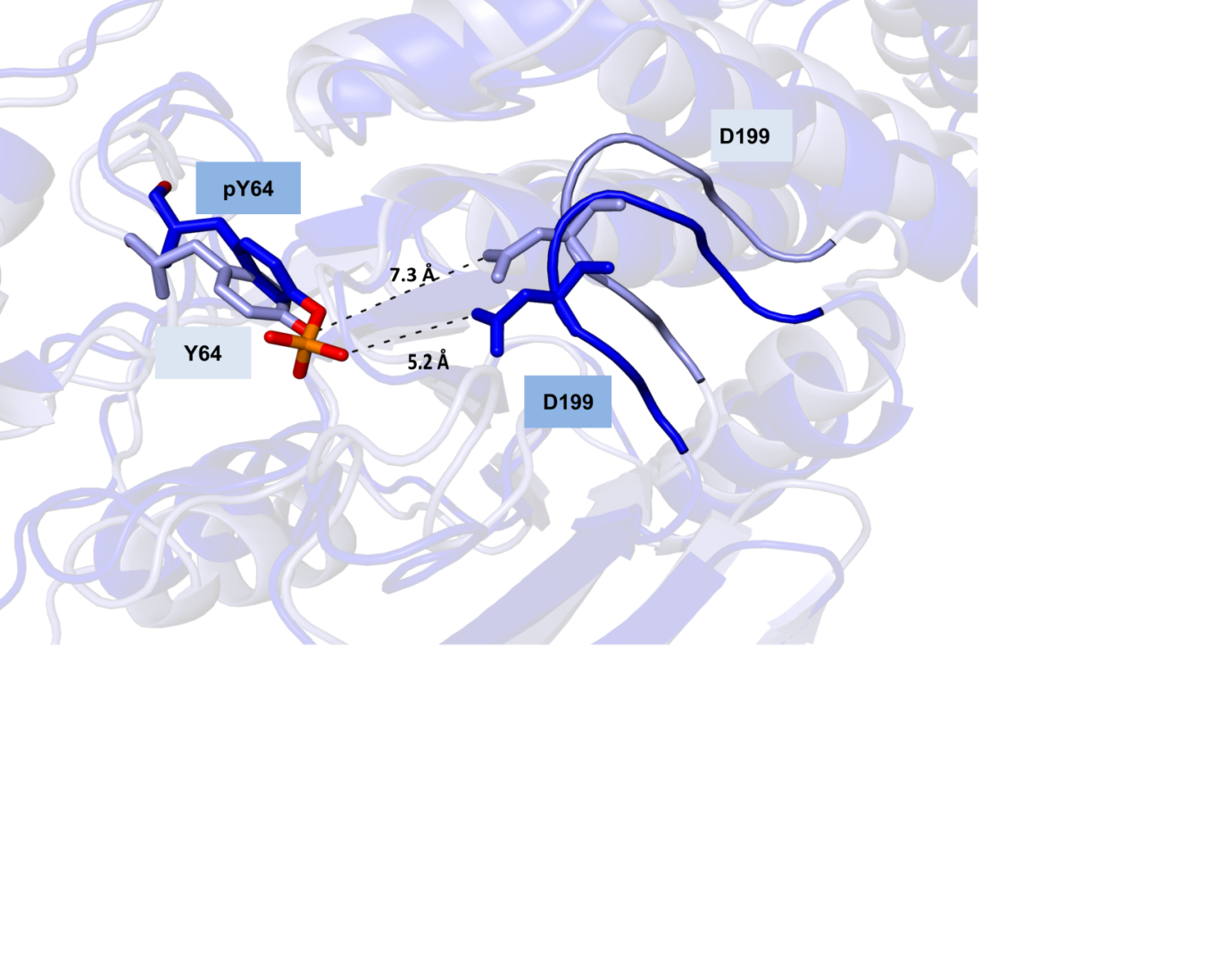

**Supplementary Figure S10. Stick representation of the interactions made by the Y64 or pY64 residues in the control and pY64 complexes respectively.** Phosphorylation of Y64 on PTP-PEST (dark blue), brings the residue pY64 closer to the WPD loop as evidenced by the shorter distance between pY64 and D199, when compared to Y64 and D199 of the control complex (light blue).

**Supplementary Figure S11**

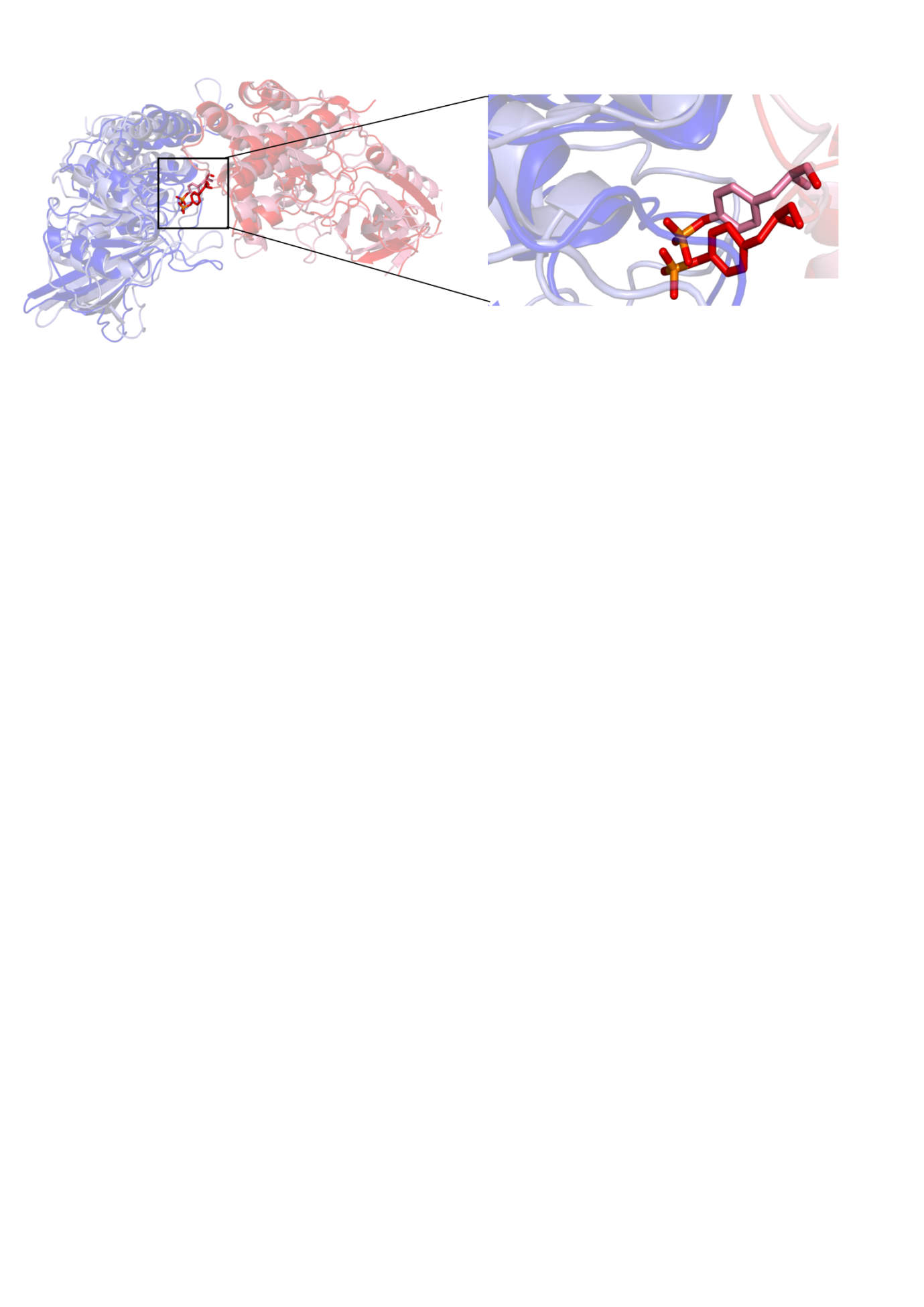

**Supplementary Figure S11. Comparison of interfaces between the control and pY64 complexes.** Phosphorylation of Y64 on PTP-PEST, leads to a deeper insertion of the residue pY232 on AMPK in the pY64 complex (red) as compared to the control complex (pink).

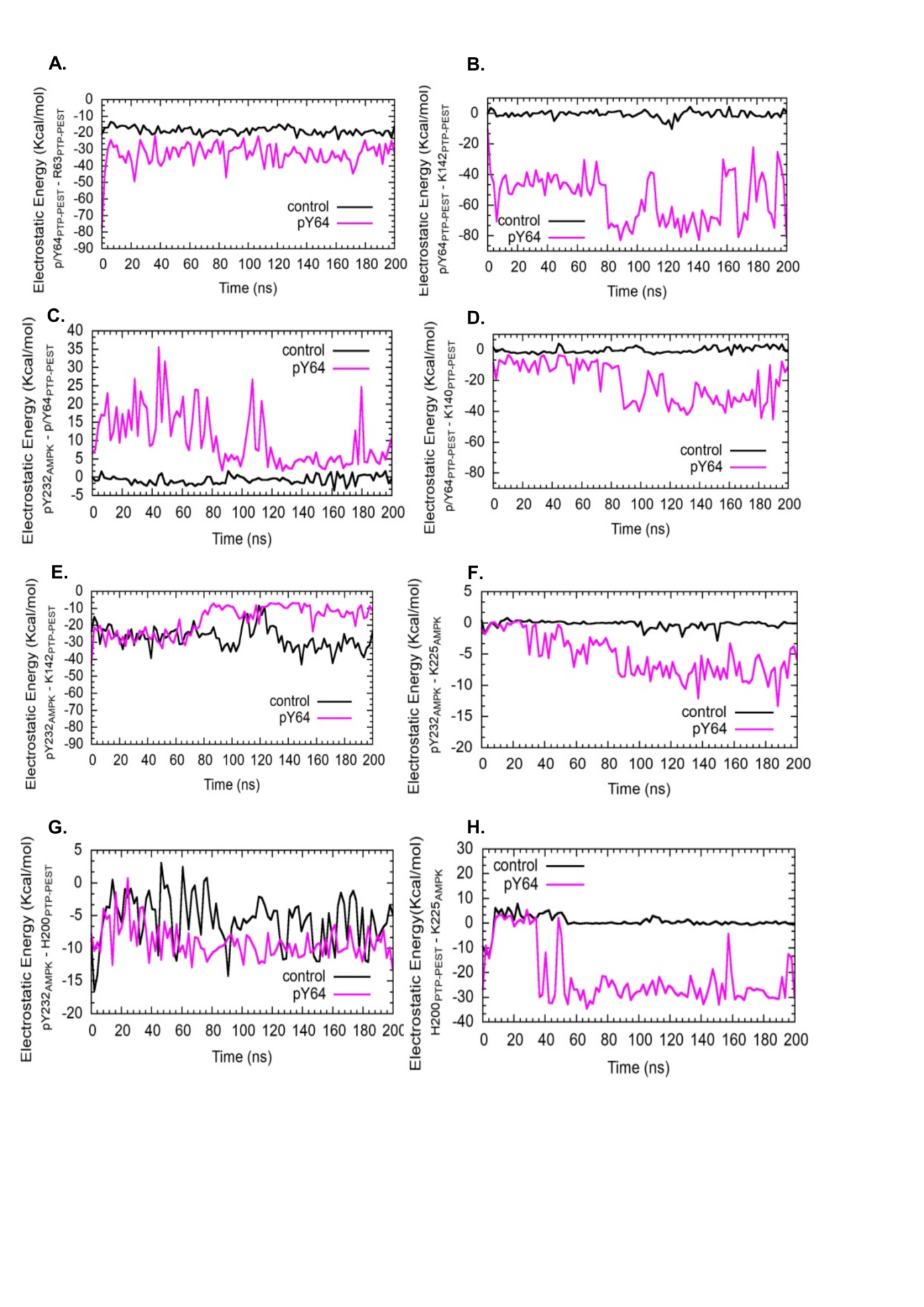
**Supplementary Figure S12. Effect of phosphorylation of Y64 on electrostatic energy of pY232 and positively charged residues in the vicinity.** Electrostatic energy of (A) pY64 of PTP-PEST in the pY64 complex or Y64 of the control complex, with R63 on PTP-PEST. Electrostatic energy of (B) pY64 of PTP-PEST in the pY64 complex or Y64 of the control complex, with K142 on PTP-PEST. Electrostatic energy of (C) pY232 of AMPK from the control and pY64 complexes with Y64 or pY64 of PTP-PEST, respectively. Electrostatic energy of (D) pY64 of PTP-PEST in the pY64 complex or Y64 of the control complex, with K140 on PTP-PEST. Electrostatic energy of	(E) pY232 of AMPK from the control and pY64 complexes with K142 of PTP-PEST. Electrostatic energy of	(F) pY232 of AMPK from the control and pY64 complexes with K225 on AMPK. Electrostatic energy of	(G) pY232 of AMPK from the control and pY64 complexes with H200 on PTP-PEST. The plots for the control and pY64 PTP-PEST—AMPK complexes are in black and pink, respectively.

**Supplementary Figure S12**

**Supplementary Figure S13**

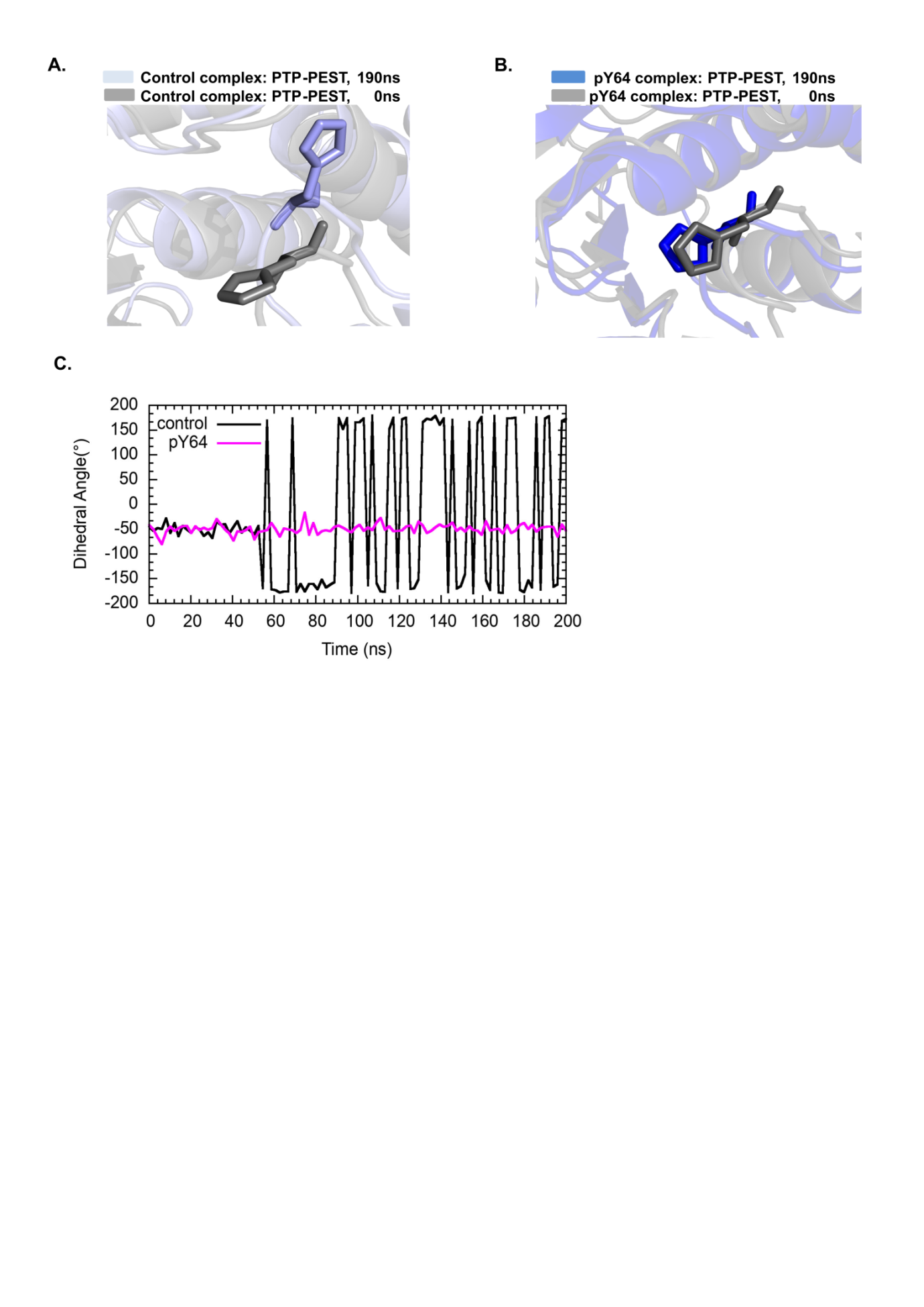

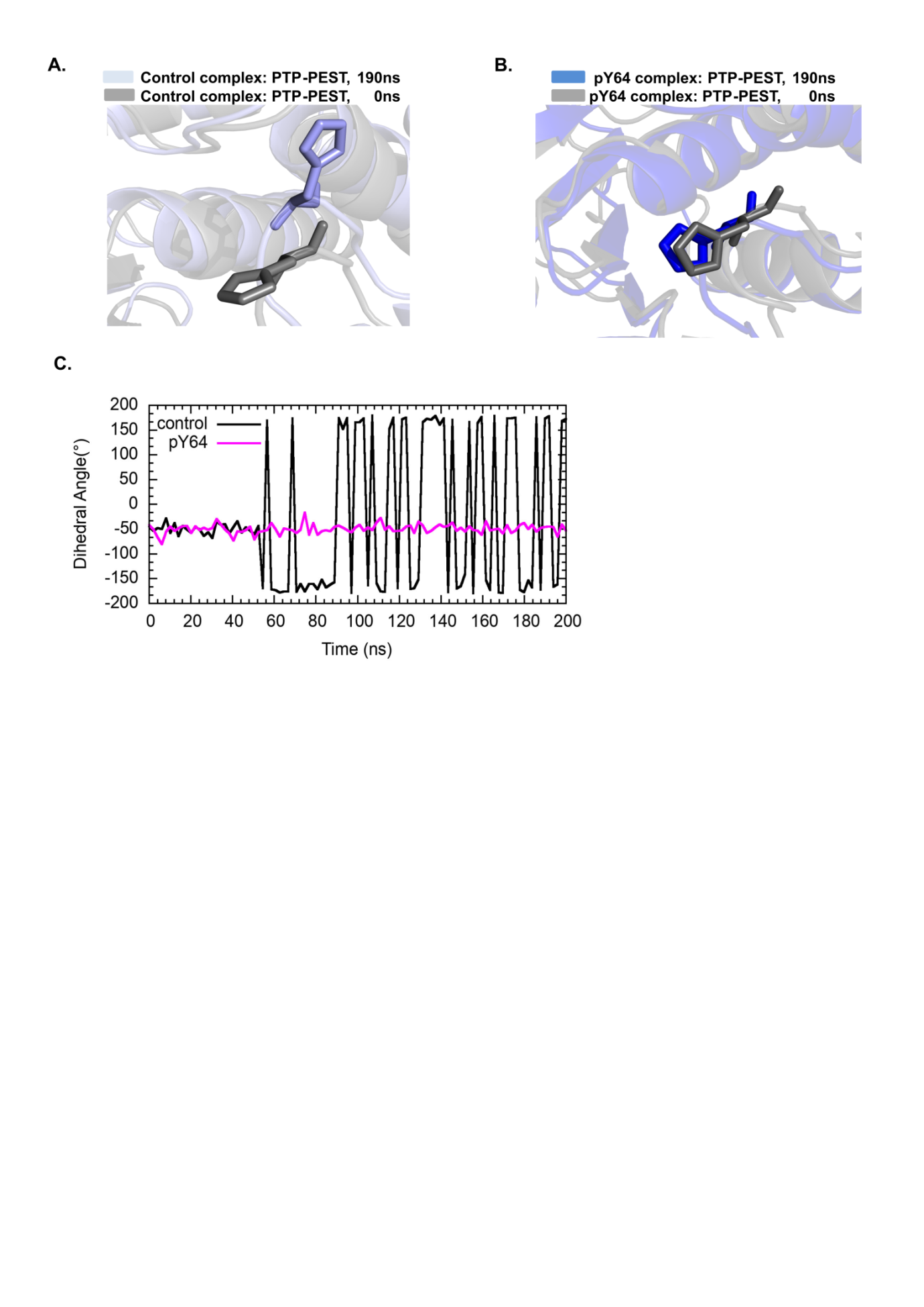
**A. B.**

**C.**

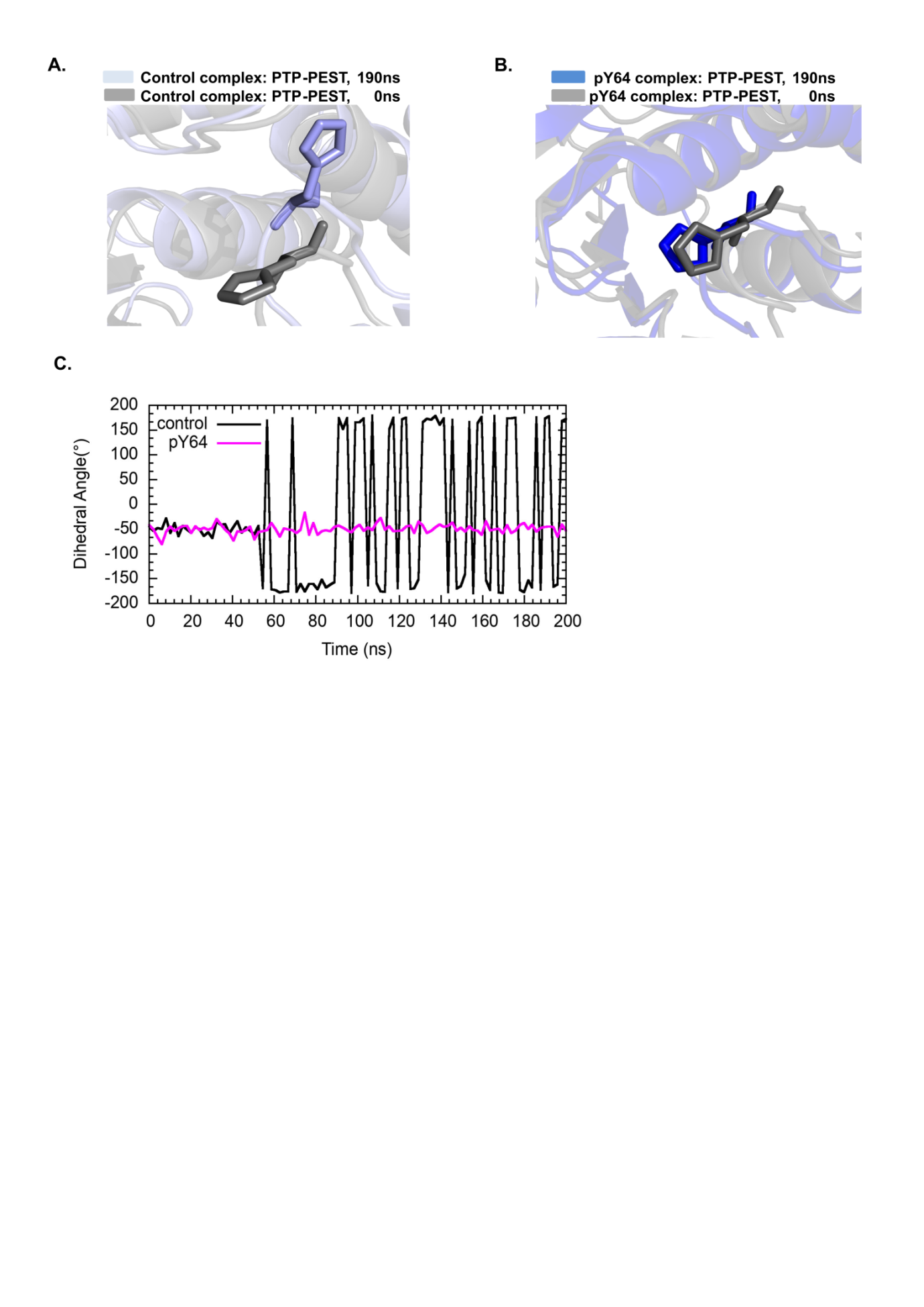

**Supplementary Figure S13. H200 Dynamics upon pY64 phosphorylation on PTP-PEST**. (A) Difference in H200 conformation between 0ns (gray) and 190ns (light blue) in the control complex (B) No change in H200 conformation in the pY64 complex at 0ns (gray) and 190ns (blue) (C) Time evolution of dihedral angle made by N-CA-CB-CG in H200, represented by black for the control and pink for the pY64 PTP-PEST—AMPK complexes.

**Supplementary Figure S14**

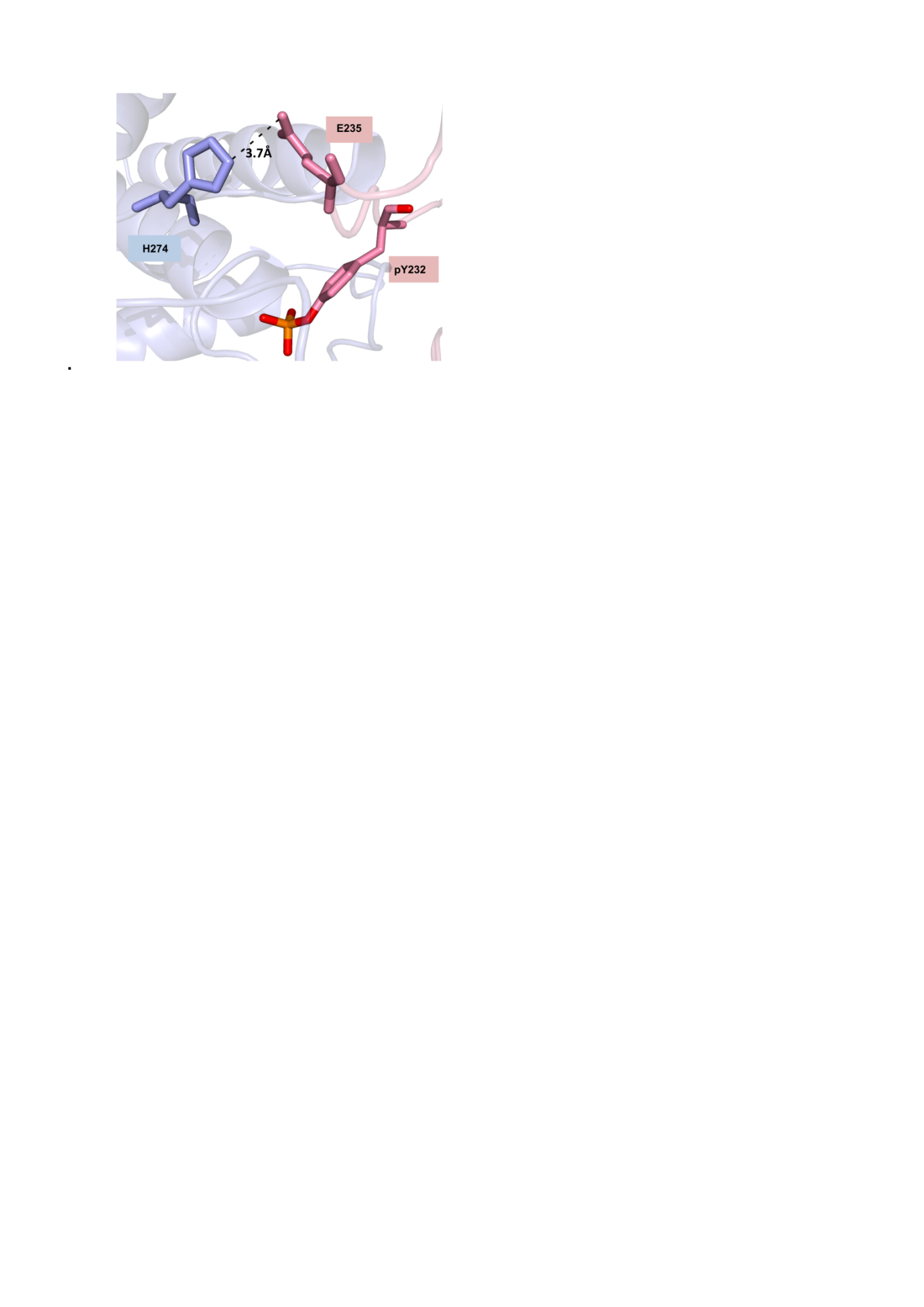

**Supplementary Figure S14. Interface of the docked complex between between PTP-PEST and AMPK phosphorylated at Y232.** The residue H274 of PTP-PEST (light blue) is within 4 Å of E235 (pink) of AMPK. The dashed line represents the distance between the two residues.

**Supplementary Tables**

**Table S1. Prediction of putative tyrosine phosphorylation sites on human AMPKα2.** (A) Predicted phosphorylation scores from the database NetPhos3.1 for tyrosines on AMPKα2. (B) Potential tyrosine phosphorylation sites and their likely tyrosine kinases, predicted from GPS 3.0.

| **Tyrosine** | **Kinases acting at site** |
| --- | --- |
| 16 | Ack |
| 80 | FGFR2, FGFR4 |
| 95 | EphA3, RON, ITK |
| 104 | FLT3 |
| 129 | FAK, EphA2, FGFR3 |
| 179 | Abl, Met, ERBB2, EphA2, EphB2, RKB, HCK |
| 190 | Axl, DDR, Anl, DDR2, EphB1, PYK2, TIE2 |
| 205 | Axl |
| 236 | InsR, EGFR, FGFR4 |
| 275 | Abl, BEGFR,EphB1, EphB2, FGFR2 |
| 283 | Syk, EphB2, SRC, YES |
| 312 | DDR, Tec, DDR2 EphB5, BTK, TEC,FLT1 |
| 324 | Ack, JAK3 |
| 341 | Trk, TYK2, MET, TEC, TXK, TRKB |
| 413 | CSK, EphA8, EphB5, JAK2, BTK |
| 436 | Ack, TNK2 |
| 450 | EphA2, JAK1, JAK2, TYK2, MET, KIT, TDGFR2, ZAP70, TXK |
| 465 | Eph, PDGFR |

| **Tyrosine Site** | **Phosphorylation score** |
| --- | --- |
| 16 | 0.387 |
| 80 | 0.427 |
| 95 | 0.409 |
| **104** | **0.629** |
| 129 | 0.415 |
| **179** | **0.639** |
| 190 | 0.39 |
| 205 | 0.397 |
| 232 | 0.433 |
| 236 | 0.448 |
| 275 | 0.388 |
| **283** | **0.962** |
| 312 | 0.421 |
| 324 | 0.353 |
| 341 | 0.402 |
| 413 | 0.66 |
| 420 | 0.4477 |
| 436 | 0.422 |
| 450 | 0.517 |
| 458 | 0.3777 |
| 465 | 0.393 |

**(A)	(B)**

**Table S2.** Ranking of docked complexes for PTP-PEST—AMPK(pY104) indicating the balanced weighted score as evaluated by [ClusPro](https://cluspro.bu.edu/login.php).

| **Cluster** | **Members** | **Representative** | **Weighted Score** |
| --- | --- | --- | --- |
| 0 | 135 | Center | -1005.9 |
|  |  | Lowest Energy | -1141.1 |
| 1 | 89 | Center | -911.8 |
|  |  | Lowest Energy | -1046.7 |
| 2 | 73 | Center | -942.6 |
|  |  | Lowest Energy | -1063.3 |
| 3 | 60 | Center | -1027.7 |
|  |  | Lowest Energy | -1027.7 |
| 4 | 54 | Center | -838.3 |
|  |  | Lowest Energy | -1072.2 |
| 5 | 48 | Center | -981.9 |
|  |  | Lowest Energy | -981.9 |
| 6 | 46 | Center | -850.7 |
|  |  | Lowest Energy | -932.5 |
| 7 | 46 | Center | -827.2 |
|  |  | Lowest Energy | -1012.5 |
| 8 | 45 | Center | -890.2 |
|  |  | Lowest Energy | -1005.8 |
| 9 | 40 | Center | -851.3 |
|  |  | Lowest Energy | -960.7 |

**Table S3.** Ranking of docked complexes for PTP-PEST—AMPK(pY283) indicating the balanced weighted score as evaluated by [ClusPro](https://cluspro.bu.edu/login.php).

| **Cluster** | **Members** | **Representative** | **Weighted Score** |
| --- | --- | --- | --- |
| 0 | 146 | Center | -921.8 |
|  |  | Lowest Energy | -1102.9 |
| 1 | 99 | Center | -978 |
|  |  | Lowest Energy | -1089.9 |
| 2 | 98 | Center | -916 |
|  |  | Lowest Energy | -1091.4 |
| 3 | 83 | Center | -934.4 |
|  |  | Lowest Energy | -1082.6 |
| 4 | 83 | Center | -1030.2 |
|  |  | Lowest Energy | -1142.6 |
| 5 | 74 | Center | -915.6 |
|  |  | Lowest Energy | -1022.2 |
| 6 | 49 | Center | -958.5 |
|  |  | Lowest Energy | -1007.8 |
| 7 | 47 | Center | -996 |
|  |  | Lowest Energy | -1094.2 |
| 8 | 46 | Center | -934.3 |
|  |  | Lowest Energy | -1061.6 |
| 9 | 44 | Center | -986.7 |
|  |  | Lowest Energy | -1147.4 |

**Table S4.** Ranking of docked complexes for PTP-PEST—AMPK(pY232) indicating the balanced weighted score as evaluated by [ClusPro](https://cluspro.bu.edu/login.php)

| **Cluster** | **Members** | **Representative** | **Weighted Score** |
| --- | --- | --- | --- |
| 0 | 212 | Center | -1018.3 |
|  |  | Lowest Energy | -1236.2 |
| 1 | 102 | Center | -970.7 |
|  |  | Lowest Energy | -1213.4 |
| 2 | 100 | Center | -961.1 |
|  |  | Lowest Energy | -1082.4 |
| 3 | 94 | Center | -996.6 |
|  |  | Lowest Energy | -1158.6 |
| 4 | 66 | Center | -1045 |
|  |  | Lowest Energy | -1204 |
| 5 | 62 | Center | -989.6 |
|  |  | Lowest Energy | -1242 |
| 6 | 51 | Center | -950.7 |
|  |  | Lowest Energy | -1209 |
| 7 | 50 | Center | -970.7 |
|  |  | Lowest Energy | -1191.9 |
| 8 | 46 | Center | -951.6 |
|  |  | Lowest Energy | -1117.2 |
| 9 | 44 | Center | -974.5 |
|  |  | Lowest Energy | -1046.6 |

**Table S5.** Summary of interface interactions of the representative docked complexes

| **Model** | **Chain** | **No. of interface residues** | **Interface area (Å^2^)** | **Salt bridges** | **Hydrogen bonds** | **Non- bonded contacts** | **FoldX Interaction Energy (Kcal/mol)** |
| --- | --- | --- | --- | --- | --- | --- | --- |
| **PTP-PEST**—**AMPK(pY104)** | **A** | 30 | 1423 | 4 | 23 | 217 | -13.29 |
|  | **B** | 27 | 1447 |  |  |  |  |
| **PTP-PEST**—**AMPK(pY232)** | **A** | 30 | 1468 | 8 | 39 | 290 | -20.17 |
|  | **B** | 36 | 1395 |  |  |  |  |
| **PTP-PEST**—**AMPK(pY283)** | **A** | 25 | 971 | 3 | 16 | 169 | -17.95 |
|  | **B** | 19 | 1065 |  |  |  |  |
